# Expanded CAG/CTG Repeats Resist Gene Silencing Mediated by Targeted Epigenome Editing

**DOI:** 10.1101/368480

**Authors:** Bin Yang, Alicia C. Borgeaud, Lorène Aeschbach, Oscar Rodríguez-Lima, Gustavo A. Ruiz Buendía, Cinzia Cinesi, Tuncay Baubec, Vincent Dion

## Abstract

Expanded CAG/CTG repeat disorders affect over 1 in 2500 individuals worldwide. Potential therapeutic avenues include gene silencing and modulation of repeat instability. However, there are major mechanistic gaps in our understanding of these processes, which prevent the rational design of an efficient treatment. To address this, we developed a novel system, ParB/ANCHOR-mediated Inducible Targeting (PInT), in which any protein can be recruited at will to a GFP reporter containing an expanded CAG/CTG repeat. Using PInT, we found no evidence that the histone deacetylase HDAC5 or the DNA methyltransferase DNMT1 modulate repeat instability upon targeting to the expanded repeat, suggesting that their effect is independent of local chromatin structure. Unexpectedly, we found that expanded CAG/CTG repeats reduce the effectiveness of gene silencing mediated by HDAC5 or DNMT1 targeting. The repeat-length effect in gene silencing by HDAC5 was abolished by a small molecule inhibitor of HDAC3. Our results have important implications on the design of epigenome editing approaches for expanded CAG/CTG repeat disorders. PInT is a versatile synthetic system to study the effect of any sequence of interest on epigenome editing.

## Introduction

Huntington’s disease and myotonic dystrophy type 1 are the two most common expanded CAG/CTG repeat diseases, accounting for over 1 in 2500 individuals world-wide [1–4]. Neuromuscular and neurodegenerative phenotypes are caused by the expression of an expanded allele that generates toxic RNAs and/or peptides, which affect gene expression, splicing, and protein aggregation *in trans* [5–8]. These mechanisms are thought to be worsened by somatic expansion of the expanded allele, which occurs in afflicted individuals over their lifetime [8]. Indeed, longer repeats cause more severe phenotypes [9,10]. Currently, there is no cure for these diseases, but modulating somatic expansion or inducing contractions are being explored as therapeutic approaches [8,11].

Expanded CAG/CTG repeats affect gene expression of the gene they reside in as well as neighbouring ones [12]. These changes in expression are associated with gains in heterochromatin marks, including histone H3 lysine 9 methylation (H3K9me), HP1 binding and CpG methylation, as well as loss of euchromatic markers, such as CTCF binding and H3 tail acetylation [13–19]. However, CAG/CTG repeat expansion does not appear to alter three-dimensional chromatin conformation [20]. Although the heterochromatic-like state reduces the expression of the mutant allele, it does not completely abolish it [12]. Furthermore, the remaining transcription through the repeat tract would be expected to support repeat instability [21]. Thus, targeting the expanded allele for silencing may provide much needed symptomatic relief.

Here we asked whether epigenome editing could be harnessed to modulate gene expression and CAG/CTG repeat instability. To this end, we developed a synthetic method that enables the targeting of any peptide to a sequence of choice embedded within the intron of a fluorescent reporter. We named the system ParB/ANCHOR-mediated induced targeting (PInT). To test our system, we inserted CAG/CTG repeats within the reporter cassette such that we could monitor both their instability as well as their effect on gene expression. Using PInT, we clarified the role of two heterochromatin proteins, histone deacetylase 5 (HDAC5) and DNA methyltransferase 1 (DNMT1) in modulating repeat instability through their local recruitment. Moreover, we show, unexpectedly, that gene silencing efficiency brought about by the targeting of either HDAC5 or DNMT1is reduced at expanded repeats compared to shorter ones. We further implicate the catalytic activity of Histone deacetylase 3 (HDAC3) in helping expanded CAG/CTG repeats resist gene silencing. Our results provide novel mechanistic insights into how HDAC5 and DNMT1 impact repeat instability and uncover an unexpected effect of repeat expansion on epigenome editing.

## Results

### ParB/ANCHOR-mediated induced targeting (PInT)

We designed PInT (Fig. 1) to be modular and highly controllable. It contains a GFP mini gene that harbours two GFP exons flanking the intron of the rat *Pem1* gene [11,22]. A doxycycline-inducible promoter drives the expression of the reporter. This cassette is always inserted at the same genomic location as a single copy integrant on chromosome 12 of T-Rex Flp-In HEK293 cells [20]. Within the intron, we inserted a 1029 bp non-repetitive sequence, *INT*, that contains four binding sites for dimers of the *Burkholderia cenocepacia* ParB protein [23]. Once bound to *INT*, ParB oligomerizes in a sequence-independent manner, recruiting up to 200 ParB molecules [24]. This ParB/ANCHOR system was first used in live yeast cells to visualize double-strand break repair [23]. More recently, it has been used to monitor the mobility of a genomic locus upon activation of transcription [25,26] and to visualize viral replication [27] and the translocation of the HIV pre-integration complex to the nucleus [28] of live mammalian cells. We made the system inducible by fusing ParB to a domain of the *A. thaliana* protein ABSCISIC ACID INSENSITIVE 1 (ABI), which dimerizes with a domain of PYRABACTIN RESISTANCE1-LIKE 1 (PYL) upon addition of abscisic acid (ABA) to the culture medium [29]. ABA is a plant hormone that is not toxic to human cells, making its use especially convenient. Within 319bp of the *INT* sequence, there is a cloning site that can be used to insert any DNA motif (Fig. 1). Fusing any protein of interest to PYL allows for full temporal control over the recruitment of a protein of interest near a DNA sequence of choice. In this case we used a CAG/CTG repeat that was either in the non-pathogenic (16 repeats) or pathogenic (≥59 repeats) range. The CAG/CTG repeats affect splicing of the reporter in a length-dependent manner, with longer repeats leading to more robust insertion of an alternative CAG exon that includes 38 nucleotides downstream of the CAG, creating a frameshift [30]. Thus, we can monitor repeat size as well as changes in gene expression upon targeting any protein of choice near a CAG/CTG repeat of various sizes.

**Fig. 1:**
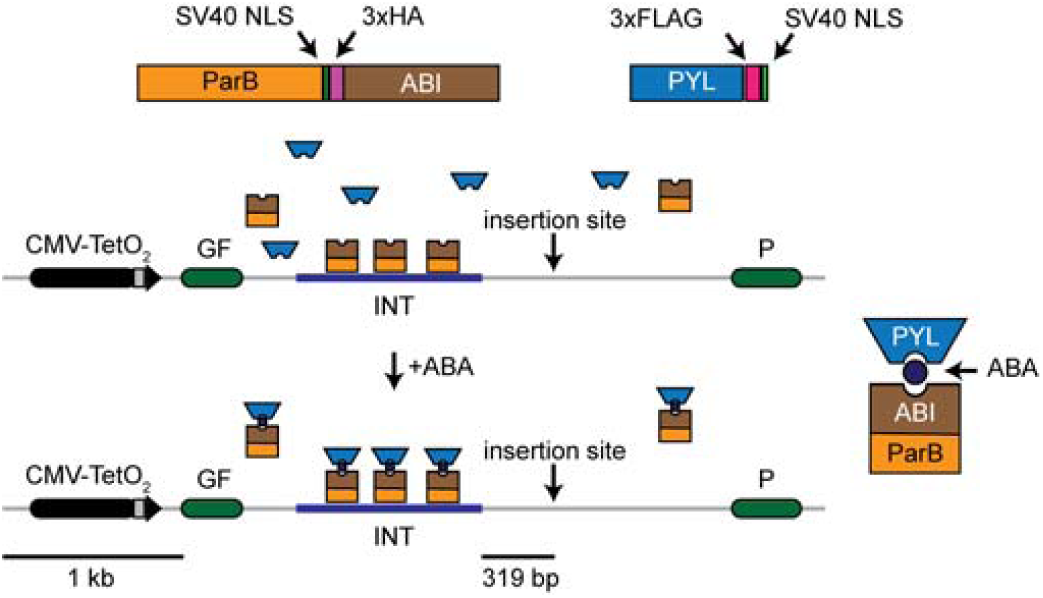
Schematic of PInT. The GFP reporter is driven by an inducible Tet-ON promoter. It contains an intron harbouring an INT sequence, which mediates the recruitment and oligomerization of ParB. We fused ParB to ABI, a plant protein that binds PYL only in the presence of abscisic acid (ABA). The PYL construct contains 3 tandem FLAG tags and the ParB-ABI fusion includes 3 tandem HA tags. They both contain SV40 nuclear localization signals. Fusing PYL to any protein of interest leads to its inducible recruitment 319bp away from a cloning site that can be used to insert a sequence of choice. Here we chose expanded CAG/CTG repeats to test the system.

First, we determined whether the components of PInT affect the expression of the GFP reporter. We tested whether ABA changed GFP expression in GFP(CAG)_0_ cells [22]. These cells carry the GFP mini gene without the *INT* sequence and no repeat in the intron (see Table S1 and Fig. S1 - for details about cell line construction). We found that treatment with up to 500 µM of ABA, which induces the dimerization between PYL and ABI [29], had no effect on GFP expression (Fig. S2AB). We also transiently transfected GFP(CAG)_0_ cells with plasmids expressing the ParB-ABI fusion. This had no detectable effect on GFP expression (Fig. S2C). We next inserted the *INT* sequence inside the *Pem1* intron and integrated this construct using site-directed recombination, generating GFP-INT cells. These cells do not express ParB-ABI. We found that the insertion of the *INT* sequence had little, if any, discernible effect on GFP expression (Fig. S2D). We conclude that individually the components of PInT do not interfere with GFP expression.

We then stably integrated the ParB-ABI fusion into GFP-INT cells to generate GFP-INT-B cells. We found a decrease in GFP expression that correlated with higher levels of ParB-ABI (Fig. S2EFG), suggesting that the binding of ParB-ABI has a predictable effect on the expression of the GFP reporter. Because of this, we integrated ParB-ABI early in the cell line construction pipeline such that all the cell lines presented here express the same amount of ParB-ABI (Fig. S1, Fig. S3, Table S1).

Next, we determined the efficiency of ABA-mediated targeting PYL to the INT sequence and the consequences on GFP expression and repeat instability. We used nB-Y cells, which contain the GFP mini gene with the *INT* sequence, stably express both ParB-ABI (B) and PYL (Y), and contain *n* CAG repeats. In this case, we used either 16 CAG repeats, which is in the non-pathogenic range, or an expanded repeat of 91 triplets (Fig. 2A). Using chromatin immunoprecipitation followed by qPCR (ChIP-qPCR), we found that only 0.02% ± 0.02% and 0.1% ± 0.04% of the input *INT* DNA could be precipitated when we treated the cells with the solvent, DMSO, alone for 5 days in a cell line with 16 or 91 CAG repeats, respectively (Fig. 2B). By contrast, the addition of ABA dissolved in DMSO to the cell media increased the association of PYL to the *INT* locus significantly, reaching 1.9% ± 0.4% and 2.5% ± 0.3% of the input pulled down in 16B-Y or 91B-Y cells, respectively (Fig. 2B – P=0.002 and P=9×10^−5^, comparing DMSO and ABA, for 16B-Y and 91B-Y, respectively, using a one-way ANOVA). At the *ACTA1* locus, where there is no *INT*, the immunoprecipitated DNA remained below 0.04% regardless of the cell line or conditions used (Fig. 2B). These results demonstrate the inducible nature of the system and show that the efficiency of the targeting is similar regardless of repeat size (P=0.2 comparing ABA conditions in 91B-Y and 16B-Y lines using a one-way ANOVA). Importantly, PYL targeting had no effect on GFP expression as measured by flow cytometry (Fig. 2C – P= 0.87 and P= 0.76, when comparing the mean GFP intensities upon DMSO or ABA treatment in 16B-Y and 91B-Y lines, respectively, using a one-way ANOVA). Moreover, targeting PYL to expanded CAG/CTG repeats by adding ABA to the medium of 91B-Y cells for 30 days had no effect on repeat instability (Fig. 2D, Table 1, P= 0.53 comparing the number of expansions, contractions, and no change in cells treated with DMSO alone to ABA-treated cells using a χ^2^ test). We conclude that PInT works as an inducible targeting system and that PYL targeting is efficient and does not further affect gene expression or repeat instability.

**Table 1:**
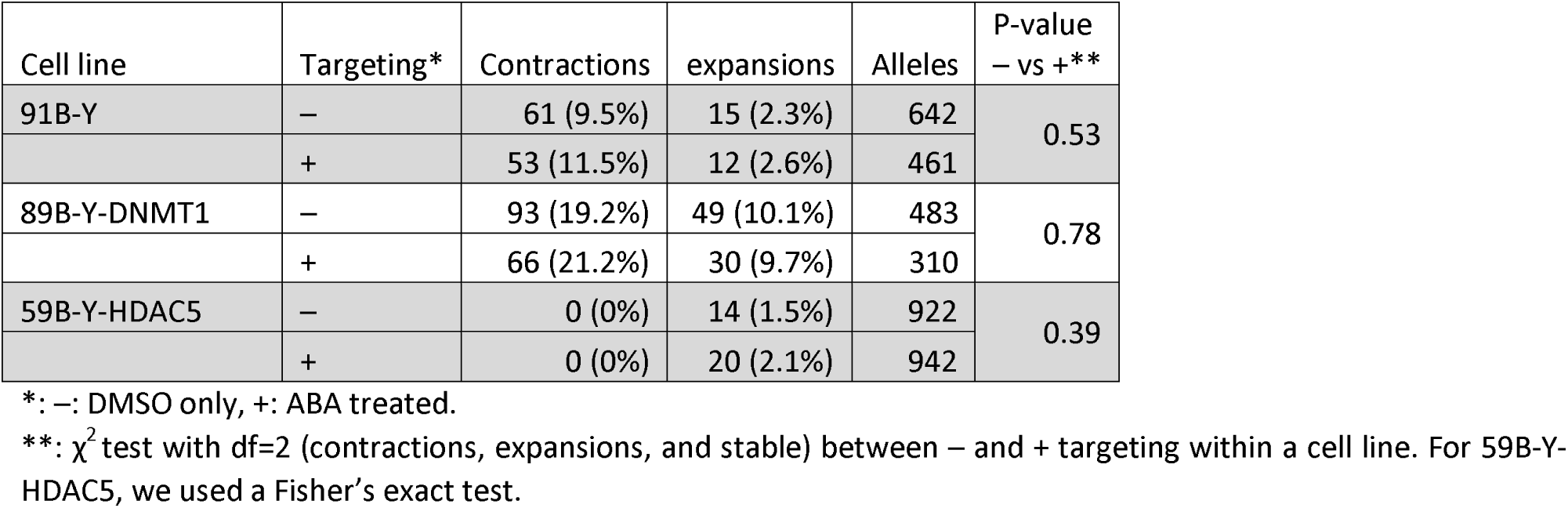
Small pool PCR quantification after 30 days of treatment with ABA or DMSO.

**Fig. 2:**
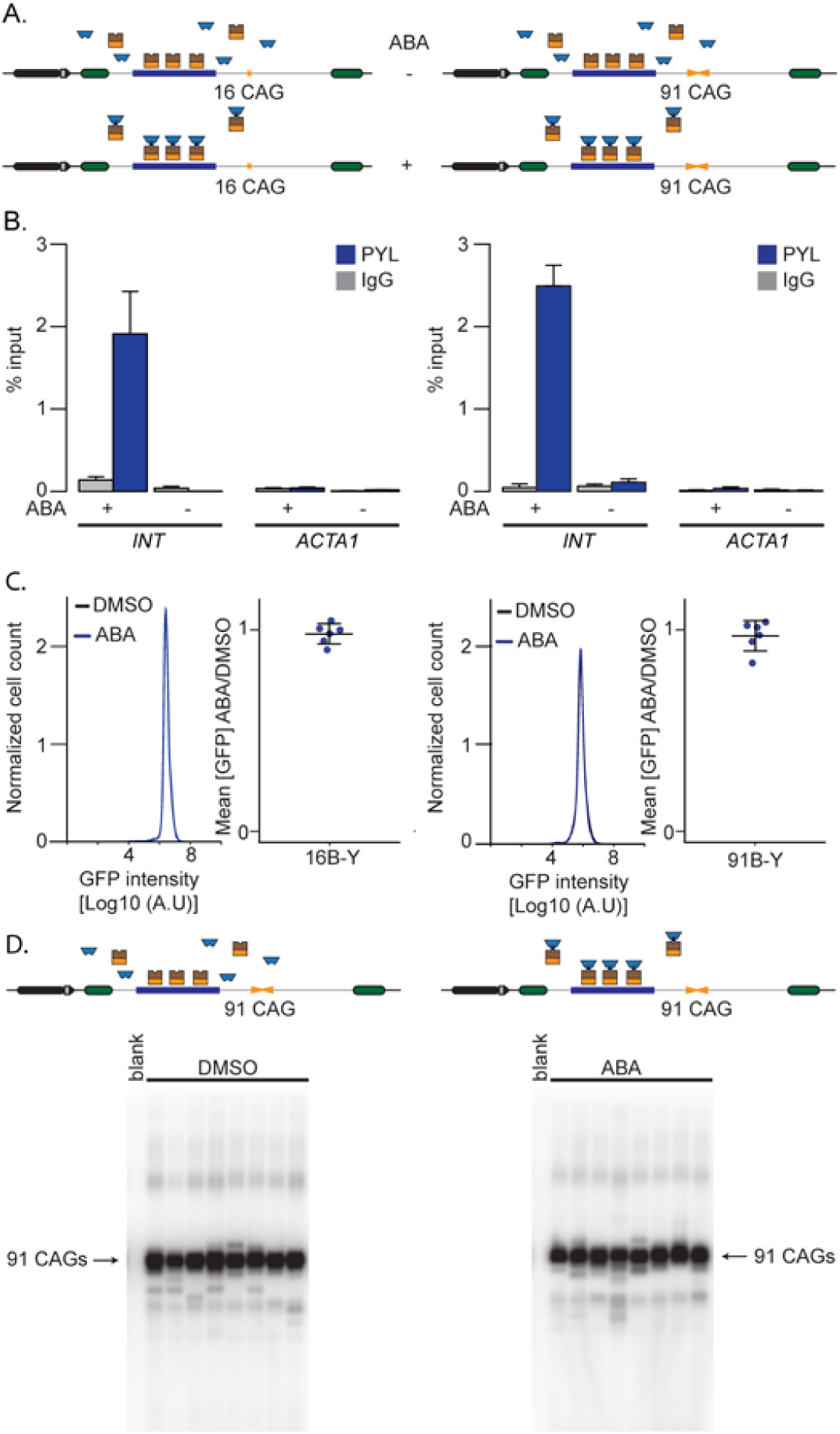
Inducible targeting of PYL at the GFP reporter. A) Schematic representation of 16B-Y (left) and 91B-Y (right) cell lines. B) ChIP-qPCR using antibodies against FLAG to pull down PYL at INT and ACTA1 in 16B-Y cells (left, N=4) and 91B-Y cells (right, N=4). The error bars represent the standard error. C) Representative flow cytometry profiles as well as quantification of the GFP expression in 16B-Y (left, N=6) and 91B-Y (right, N=6) cells. The error bars represent the standard deviation around the mean. D) Representative SP-PCR blots after 30 days of continuous culture in the presence of DMSO (left) or ABA (right) in 91B-Y cells. 1 ng of DNA/reaction used in both cases.

### Using PInT to untangle the local versus indirect roles of chromatin modifiers

Several chromatin modifiers have been implicated in CAG/CTG repeat expansion [12,15,31–36]. These studies relied on knockout or knockdown of chromatin modifiers and could not distinguish whether factors act locally at the repeat locus (i.e., *in cis*), indirectly (i.e., *in trans*), or both. PInT is designed to evaluate these possibilities. By fusing a chromatin modifier to PYL, we can induce its local recruitment and ask whether repeat instability is affected beyond any effect its overexpression has. The assumption is that overexpression levels are constant with and without ABA because it is done in the same cell line. If there is a difference in repeat instability between cells treated with ABA or those treated with DMSO alone, then we can conclude that the chromatin modifier acts locally. By contrast, a modifier that acts solely indirectly, for example by altering the transcriptome of a cell, will not show differences between ABA- and DMSO-treated cells. It is crucial to note that PInT is deliberately designed to compare non-targeted to targeted conditions within the same cell line as well as between lines with different repeat sizes (Fig. S1).

### No evidence that DNMT1 impacts repeat instability by acting *in cis*

DNMT1 maintains DNA methylation levels during replication and repair [37]. It has been implicated in preventing CAG/CTG repeat expansion in the germlines of a mouse model for spinocerebellar ataxia type 1 [31]. Heterozygous DNMT1 mice showed lower expansion in the germlines accompanied by changes in CpG methylation flanking the repeat tract in testes and ovaries. High local CpG methylation correlated with high levels of repeat instability [31], suggesting that local levels of DNA methylation promote repeat instability. Here we tested this hypothesis using PInT and targeted PYL-DNMT1 to 16 or 89 CAG/CTG repeats (Fig. 3A). The hypothesis predicts that targeting PYL-DNMT1 will increase CpG methylation near the repeat tract and thereby increase repeat expansion frequencies. ChIP-qPCR confirmed robust recruitment of PYL-DNMT1 to levels comparable to PYL alone (Fig. 3B). Indeed, enrichment rose upon addition of ABA from 0.3% ± 0.1% to 5.0% ± 0.5% and from 0.3% ± 0.2% to 6.8% ± 0.7% in 16B-Y-DNMT1 and 89B-Y-DNMT1 cells, respectively. The recruitment was statistically significant (P=8×10^−5^ and P=9×10^−5^ comparing qPCR enrichment with and without ABA in 16B-Y-DNMT1 and 89B-Y-DNMT1 lines, respectively, using a one-way ANOVA). Here again, the enrichment was not seen at the *ACTA1* locus, suggesting that it is specific to the presence of the *INT* sequence (Fig. 3B). We further determined whether targeting PYL-DNMT1 could increase levels of CpG methylation near the repeat tract. To do so, we performed bisulfite sequencing after targeting PYL-DNMT1 for 30 days. This led to changes of 10 to 20% in the levels of CpG methylation, a modest increase(Fig. 3C), which is in line with the weak de novo methyltransferase activity of DNMT1 (for example see [38,39]). Similar changes in levels of CpG methylation in Dnmt1 heterozygous ovaries and testes were seen to correlate with changes in repeat instability *in vivo* [31].

**Fig. 3:**
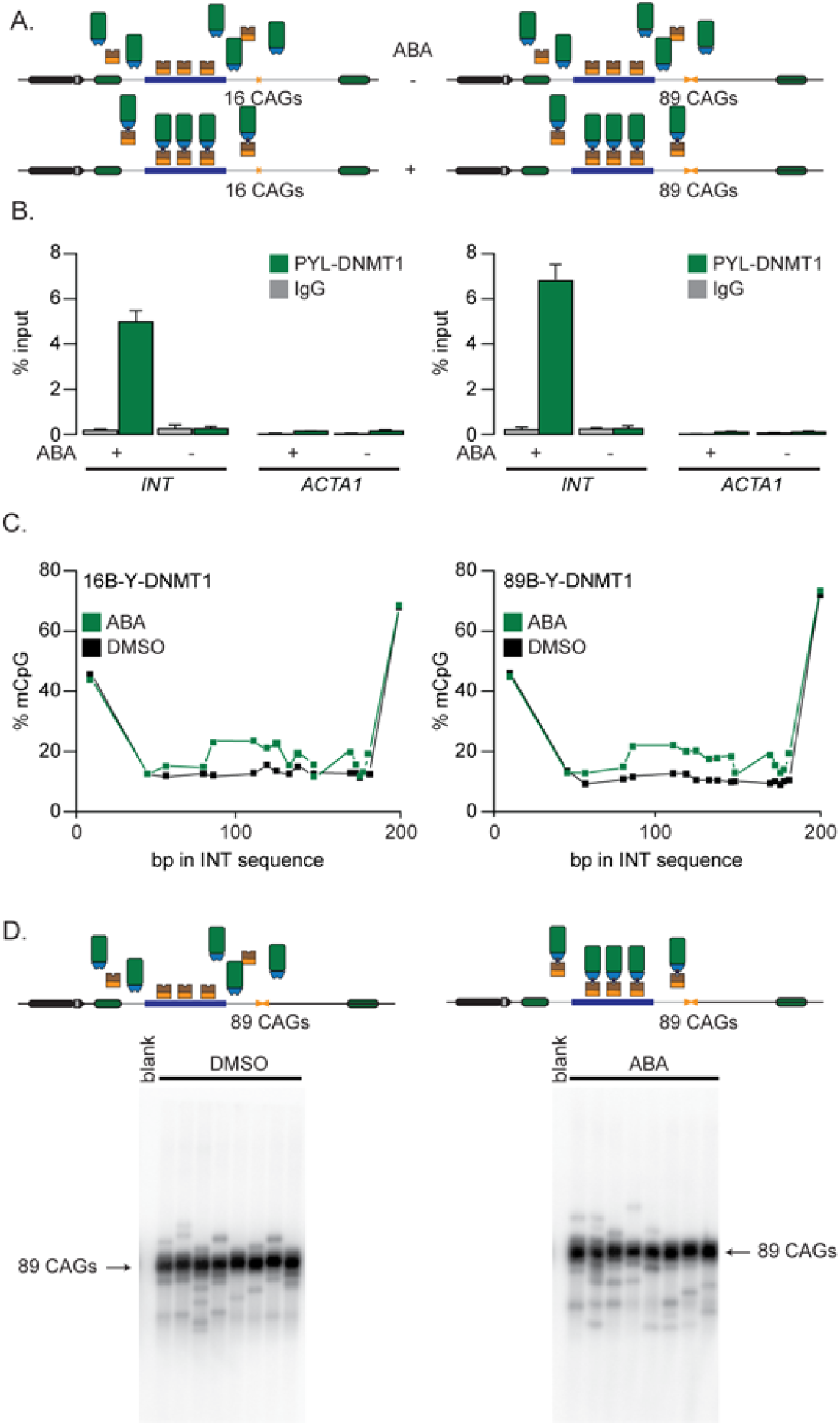
Inducible targeting of PYL-DNMT1 leads to changes in CpG methylation. A) Schematic representation of 16B-Y-DNMT1 (left) and 89B-Y-DNMT1 (right) cell lines. B) ChIP-qPCR using antibodies against FLAG to pull down PYL at INT and ACTA1 in 16B-Y-DNMT1 cells (left, N=4) and 89B-Y-DNMT1 cells (right, N=4). The error bars represent the standard error. C) Bisulfite sequencing showing the percentage of methylated CpG motifs at the INT sequence in 16B-Y-DNMT1 (left) and 89B-Y-DNMT1 (right) cells in the presence of DMSO alone (black) or ABA (green). D) Representative SP-PCR blots after 30 days of continuous culture in the presence of DMSO (left) or ABA (right) in 89B-Y-DNMT1 cells. 1 ng of DNA/reaction used in both cases.

Next, we assessed whether targeting PYL-DNMT1 promotes repeat expansion as predicted if CpG methylation increases [31]. To do so, we cultured 89B-Y-DNMT1 cells in the presence of either ABA or DMSO for 30 days, along with doxycycline to induce transcription through the repeat tract. We found no difference in the allele size distribution between the ABA and DMSO conditions as measured by small-pool PCR (Table 1, Fig. 3D - P= 0.78 using a χ^2^ test), suggesting that CpG methylation near the repeat tract is not enough to drive repeat expansion. Rather, our data argue that DNMT1 has an indirect role in CAG/CTG repeat instability.

### No evidence for a local role of HDAC5 on repeat instability

HDAC5 is a Class IIa deacetylase associated with gene silencing and heterochromatin maintenance [40,41] as well as cell proliferation [42,43]. It was found to work together with HDAC3 and the MutSβ complex to promote CAG/CTG repeat expansion in a human astrocyte cell line [33,44]. HDAC3 knock down promoted expansions without changing in histone acetylation around the repeat tract [44]. Therefore, we tested whether HDAC5 had a local role in repeat expansion by acting locally at the repeat tract. We created isogenic nB-Y-HDAC5 cells that stably express a PYL-HDAC5 fusion and contain 16 or 59 CAG repeats within the GFP reporter (Fig. 4A). We found that adding ABA to the culture medium led to an increase in pull-down efficiency of PYL-HDAC5 at the *INT* locus from 0.06% ± 0.03% to 2.2% ± 0.2% in 16B-Y-HDAC5 cells and from 0.1% ± 0.1% to 3.0% ± 0.6% in 59B-Y-HDAC5 (Fig. 4B). PYL-HDAC5 targeting reduced the levels of acetylated histone H3 (H3Ac), as measured by ChIP-qPCR (Fig. 4C - P= 1.7×10^−6^ and P= 0.039 comparing DMSO- and ABA-treated 16B-Y-HDAC5 and 59B-Y-HDAC5, respectively, using an one-way ANOVA), consistent with a functional recruitment of PYL-HDAC5 to the *INT* sequence. This was confirmed by transiently transfecting PYL-HDAC5 in GFP-INT cells, which led to slightly lower GFP expression than those expressing PYL alone (Fig. S4A). Notably, PYL recruitment led to a marginally significant decrease in H3Ac level in 16B-Y cells but the decrease was not statistically significant in 91B-Y cells (Fig. 4D - P=0.04 and P=0.28 comparing the DMSO and ABA treatments in 16B-Y and 91B-Y cells, respectively, using an one-way ANOVA). Moreover, we found no significant change in acetylation upon ABA treatment at the *ACTA1* locus in either cell lines (P > 0.09 using a one-way ANOVA comparing H3ac levels in DMSO and ABA treated cells). Interestingly, the H3ac levels at the *INT* sequence were similar between 16B-Y and 91B-Y (Fig. 4D, P=0.44 comparing DMSO treated 16B-Y and 91B-Y cells using a one-way ANOVA), suggesting that the H3ac levels are unaffected by the expansion. Our results show that targeting PYL-HDAC5 reduces the levels of acetylated H3 near the repeat tract.

**Fig. 4:**
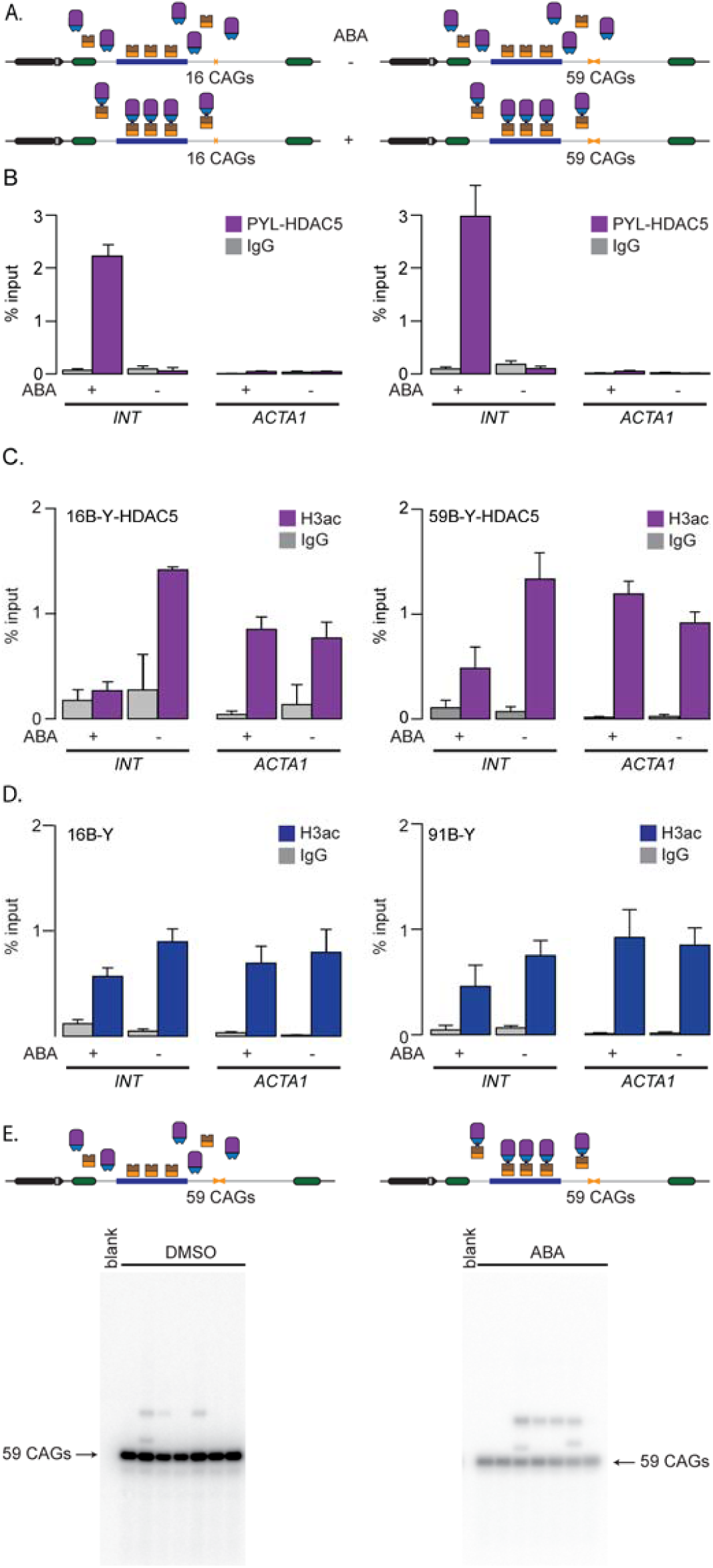
PYL-HDAC5 targeting reduces acetylation of histone H3. A) Schematic representation of 16B-Y-_HDAC5 (left) and 59B-Y-HDAC5 (right) cells. B) ChIP-qPCR using antibodies against FLAG to pull down PYL-HDAC5 at INT and ACTA1 in 16B-Y-HDAC5 cells (left, N=4) and 59B-Y-HDAC5 cells (right, N=4). The error bars represent the standard error. C) ChIP-qPCR data using a pan-acetylated H3 antibody to pull down the INT and ACTA1 loci in 16B-Y-HDAC5 (left, N=4) and 59B-Y-HDAC5 (right, N=4) cells. The error bars represent the standard error. D) ChIP-qPCR data using a pan-acetylated H3 antibody to pull down the INT and ACTA1 loci in 16B-Y (left, N=4) and 91B-Y (right, N=4) cells. The error bars represent the standard error. E) Representative SP-PCR blots after 30 days of continuous culture in the presence of DMSO (left) or ABA (right) in 59B-Y-HDAC5 cells. 1 ng of DNA/reaction used in both cases.

To monitor the local effect of PYL-HDAC5 targeting on CAG/CTG repeat instability, we cultured 59B-Y-HDAC5 cells with ABA or DMSO for 30 days. We found no difference in allele size distribution between these two treatments (Fig. 4E, Table 1 - P=0.39 using a Fisher’s exact test, comparing ABA and DMSO treated cells). Therefore, we find no evidence to support the hypothesis that HDAC5 promotes repeat expansion via local changes in protein acetylation around the repeat tract.

### Gene silencing efficiency depends on CAG/CTG repeat length

We originally designed PInT to determine whether factors work in cis for repeat instability, yet our construct also includes a GFP reporter that can be used for monitoring gene expression. This is useful to look for chromatin modifiers that can silence expanded repeats. Indeed, finding factors that, upon targeting, can silence a gene specifically when it bears an expanded allele would open doors to novel therapeutic avenues. We therefore evaluated whether DNMT1 or HDAC5 targeting could silence a reporter bearing CAG/CTG repeats, we used PInT to measure GFP expression upon ABA addition (Fig. 5A). In 16B-Y-DNMT1 cells, ABA treatment decreased GFP expression by 2.2-fold compared to DMSO treatment alone.

**Fig. 5:**
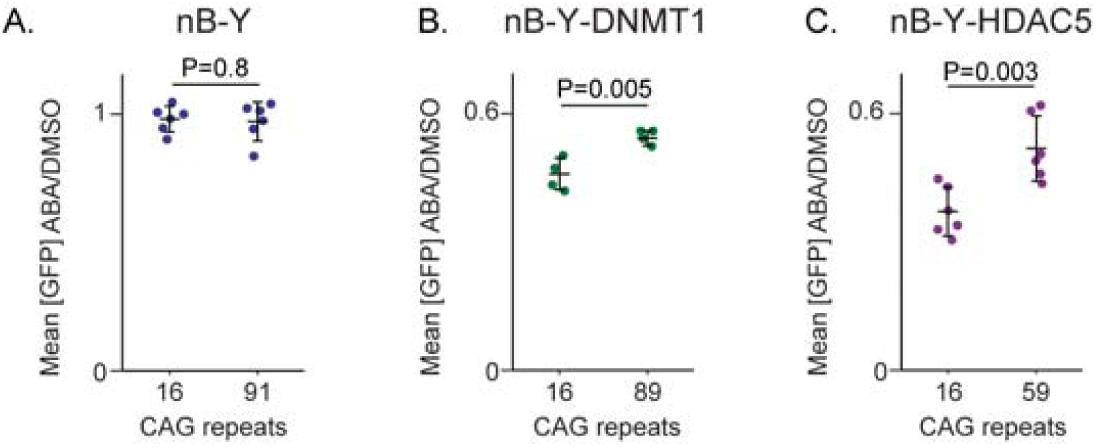
Expanded CAG/CTG repeat resist gene silencing. Mean GFP intensity ratios between ABA and DMSO alone plotted for A) 16B-Y cells (same data as in Fig. 2C – N=6), 91B-Y cells (same data as in Fig. 2C – N=6), B) 16B-Y-DNMT1 (N=4), 89B-Y-DNMT1 (N=4), C) 16B-Y-HDAC5 (N=6), and 59B-Y-HDAC5 (N=6). P-values were generated using one-way ANOVA.

Surprisingly, ABA-induced silencing was 1.8 fold compared to DMSO alone, or 16% less efficient in 89B-Y-DNMT1 than in 16B-Y-DNMT1 cells. Although relatively small, the decrease between the two lines was statistically significant (Fig. 5B, P=0.005 using a one-way ANOVA comparing the ratio of the mean GFP expression between ABA and DMSO treated cells between the two cell lines). This was not due to PYL-DNMT1 being targeted more efficiently upon ABA addition or leading to higher levels of CpG methylation around the repeat tract in 16B-Y-DNMT1 cells compared to 89B-Y-DNMT1 cells (Fig. 3BC). These results rather suggest that the presence of an expansion reduces the efficiency of PYL-DNMT1 to silence the reporter.

We next addressed whether this effect was specific to DNMT1. We added ABA to the medium of 16B-Y-HDAC5 cells for five days and found a reduction of GFP expression of 2.7-fold (Fig. 5A). This decrease in expression was significantly smaller in the context of an expanded repeat (Fig. 5C, P=0.003 comparing the decrease in expression upon ABA addition between the 16B-Y-HDAC5 and 59B-Y-HDAC5 using a one-way ANOVA). Some more mundane explanations were ruled out, including a difference in targeting efficiency of PYL-HDAC5 or changes in H3Ac levels between the cell lines (Fig. 4BC). We also tested whether the allele length-specific effect on GFP expression required the presence of the *INT* sequence. Thus, we transiently expressed PYL-HDAC5 in GFP(CAG)_0_B cells, which have no *INT* in their GFP reporter but express ParB-ABI.

Adding ABA to these cells did not affect GFP expression (Fig. S4BC), suggesting that the presence of the *INT* sequence is essential. Taken together, our results suggest that expanded CAG repeats resist gene silencing mediated by both DNMT1 and HDAC5.

### The N-terminal domain of HDAC5 mediates silencing

PInT can also be used to delineate the mechanism of gene silencing upon targeting of a chromatin modifier. To exemplify this, we sought to clarify how HDAC5 silences the reporter. Class I HDACs, like HDAC3, derive their catalytic activity *in vitro* from a conserved tyrosine residue that helps coordinate a zinc ion essential for catalysis [43]. By contrast, Class IIa enzymes, like HDAC5, have a histidine instead of tyrosine at the analogous site, which considerably lowers HDAC activity [43]. In fact, restoring the tyrosine at position 1006 of HDAC5 increases HDAC activity by over 30-fold [43]. We reasoned that if the HDAC activity was responsible for the silencing activity, the H1006Y gain-of-function mutant should lead to a more robust silencing. Moreover, the H1006A mutant, in which the HDAC activity is abolished altogether, would be expected to silence the reporter. To test these predictions, we transiently transfected PYL-HDAC5 wild-type as well as H1006Y and H1006A mutants in 40B cells, which contain the GFP-INT reporter with 40 CAGs and express ParB-ABI (Fig. 6A). Overall, the effect on silencing seen upon targeting of the wild-type PYL-HDAC5 fusion was smaller when delivered by transient transfection compared to the stable cell lines. Nevertheless, the wild-type PYL-HDAC5 significantly reduced GFP expression compared to PYL alone (P=0.00001 using a one-way ANOVA). In the same conditions, targeting PYL-HDAC5-H1006A or PYL-HDAC5-H1006Y both silenced the transgene compared to targeting PYL alone (Fig 6B; P= 0.006 and 0.002, respectively, using a one-way ANOVA), suggesting that tampering with the catalytic activity of HDAC5 does not influence silencing of our GFP reporter. Moreover, targeting PYL fused to the catalytic domain of HDAC5 did not shift GFP expression compared to PYL alone (Fig. 6B – P=0.88 using a one-way ANOVA). Rather, we find that the silencing activity was contained within the N-terminal part of HDAC5, which characterizes Class IIa enzymes. Further truncations (Fig. 6A) are consistent with a model whereby the coiled-coil domain in the N-terminal part of HDAC5, which is necessary for homo- and heterodimerization of Class IIa enzymes *in vitro* [45], contains the silencing activity (Fig. 6B). It may therefore be that this domain recruits endogenous HDACs to the locus and mediate gene silencing.

**Fig 6:**
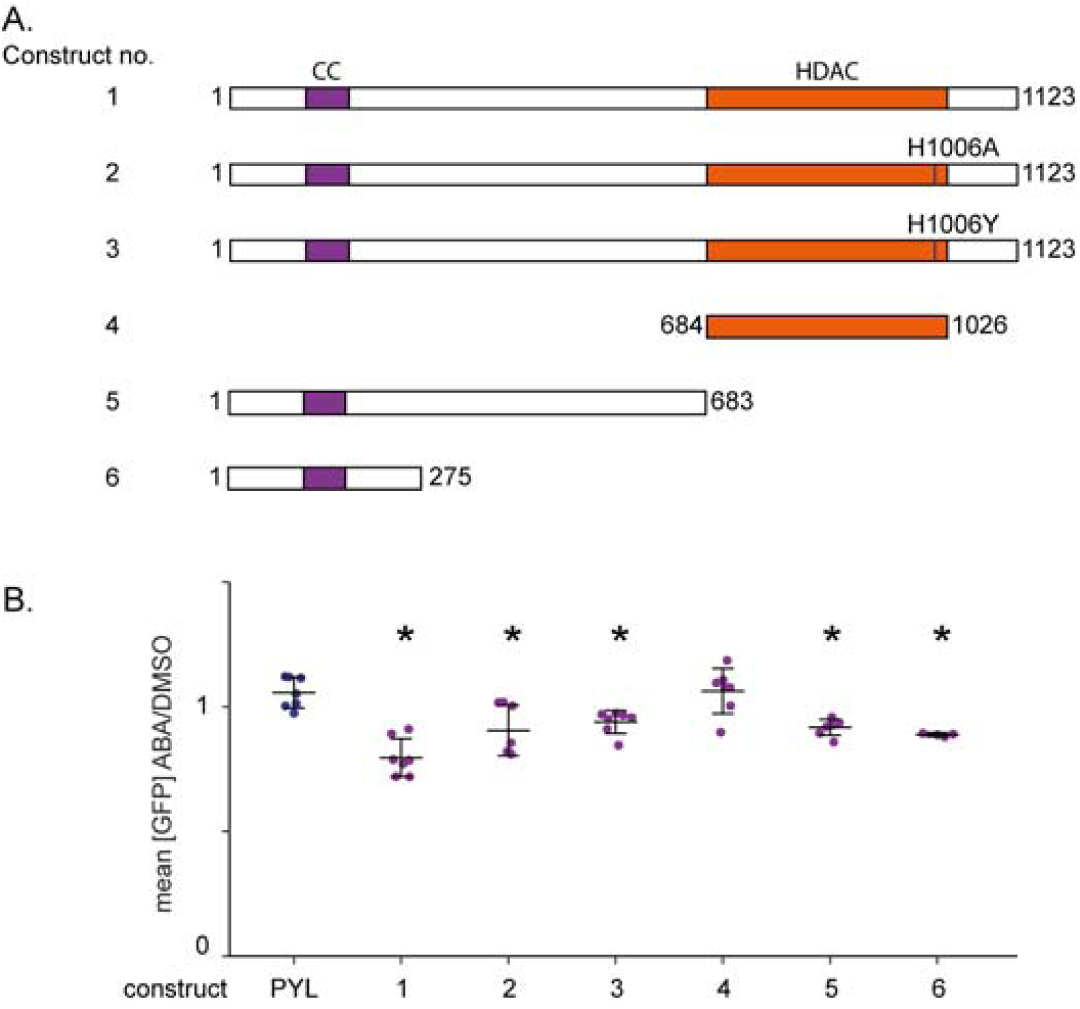
HDAC5 mediates silencing through its N-terminal. A) Mutants and truncations of HDAC5 fused to PYL. The coiled-coil (CC) domain is indicated in purple, the deacetylase domain (HDAC) in orange. B) ABA and DMSO treatments of 40B cells transiently transfected with plasmids containing the constructs shown in B. Construct 1: N=7, P = 0.00001 vs PYL; construct 2: N=7, P= 0.006 vs PYL; construct 3: N=7, P=0.001 vs PYL; construct 4: N=7, P=0.88 vs PYL; construct 5: N=7, P=0.0002 vs PYL; construct 6: N=3, P=0.0003 vs PYL. *: P≤0.01 compared to PYL targeting. The error bars show the standard deviation around the mean. P-values were generated using one-way ANOVA.

### PYL-HDAC3 targeting enhances GFP expression independently of its catalytic activity

HDAC5 is thought to mediate histone deacetylation by recruiting other HDACs, including HDAC3 [46]. Therefore, we hypothesized that PYL-HDAC3 targeting would have the same effect on GFP expression as PYL-HDAC5 targeting. To address this directly, we overexpressed PYL-HDAC3 in 40B cells without targeting (Fig. S4D). We found that there was a slight decrease in GFP expression, suggesting that the construct could silence gene expression *in trans*. Next, we generated stable nB-Y-HDAC3 cells and compared GFP intensities with and without ABA (Fig. S5A). Contrary to our initial hypothesis, we found that targeting PYL-HDAC3 in both 16B-Y-HDAC3 and 89B-Y-HDAC3 increased GFP expression by 1.5 fold (Fig. S5B, P=0.002 and P=0.02 using a one-way ANOVA comparing ABA and DMSO treatments in 16B-Y-HDAC3 and 89B-Y-HDAC3, respectively). The effect appeared to require *INT* since adding ABA to GFP(CAG)_0_B cells transiently expressing PYL-HDAC3 did not affect GFP expression (Fig. S4E). The increase in GFP expression in nB-Y-HDAC3 cells was accompanied by an efficient targeting of the PYL-HDAC3 fusion (Fig. S5C) and an increase in H3ac levels, suggesting that HDAC3 deacetylase activity is not implicated in this effect (Fig. S5D). We confirmed this by adding the HDAC3-specific small molecule inhibitor RGFP966 [47] to nB-Y-HDAC3 cells. This treatment did not affect the ability of PYL-HDAC3 of increasing GFP expression in neither 16B-Y-HDAC3 nor 89B-Y-HDAC3 lines (Fig. S5E). We therefore conclude that targeting PYL-HDAC3 increases GFP expression independently of its HDAC activity.

### HDAC3 activity is required for the repeat size-specificity upon HDAC5-mediated silencing

Next, we asked whether PInT could be used to gain insights into the mechanism of targeted epigenome editing. To do so, we sought to find enzymatic activities that can modify allele-size specific silencing brought about by PYL-HDAC5 targeting. Although HDAC3 targeting did not have the expected effect on GFP expression, evidence shows that its catalytic activity is implicated in HDAC5-mediated silencing [46]. To determine whether the catalytic role of HDAC3 was essential in HDAC5-mediated silencing, we repeated our experiments in nB-Y and nB-Y-HDAC5 lines in the presence of the HDAC3 inhibitor RGFP966 (Fig. 7). We found that this small molecule had no effect on GFP expression upon PYL targeting (Fig. 7A). However, it abolished the allele-length specificity of PYL-HDAC5 targeting, leading to a silencing efficiency of 2.4- and 2.5-fold in 16B-Y-HDAC5 and 59B-Y-HDAC5, respectively (Fig. 7B, P= 0.78 using a one-way ANOVA). This contrasts with the RGFP966-free conditions where targeting PYL-HDAC5 more effectively silenced the non-pathogenic-sized allele (Fig. 5C). These results suggest expanded CAG/CTG repeats impede PYL-HDAC5-mediated silencing via the catalytic activity of HDAC3.

**Fig. 7:**
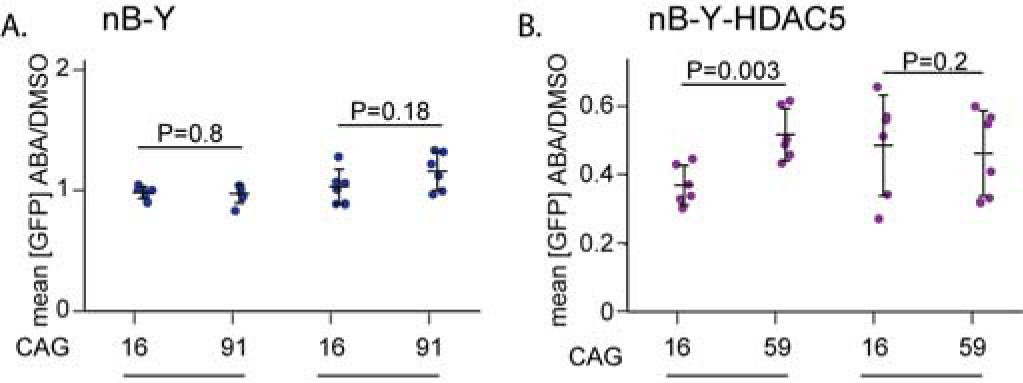
Allele-size specificity of HDAC5-mediated gene silencing requires HDAC3 activity. Quantification of GFP intensity upon targeting in the presence of 10 µM of RGFP966 or untreated A) 16B-Y (N=6 for each condition) and 91B-Y cells (N=6 for each condition) and B) 16B-Y-HDAC5 (N=6 for each condition) and 59B-Y-HDAC5 cells (N=6 for each condition). Note that the data for the untreated cells are the same as in Fig. 5. P-values were generated using one-way ANOVA.

## Discussion

Chromatin structure impinges on every DNA-based transaction, from replication and DNA repair to transcription. Consequently, epigenome editing is being harnessed to understand basic molecular mechanisms of pathogenesis and for the development of novel therapeutic approaches [48]. Epigenome editing is now most commonly carried out by fusing a chromatin modifying peptide to a catalytically dead Cas9 (dCas9). These dCas9-based approaches are highly versatile and have been used successfully to modify disease phenotypes in cells and *in vivo* [49,50]. PInT is meant to complement dCas9-based approaches. Indeed, PInT offers two advantages that we have exploited here. First, it can concentrate a large number of molecules at a target site [24], independently of chromatin context [23]. It is less practical with dCas9 to recruit equivalent numbers of molecules at a specific locus. Achieving the recruitment of a similar number of molecules at a single locus would require the use of multimerization domains, e.g., SunTag [51], in addition to multiple sgRNAs. Second, targeting a chromatin modifying peptide to different loci can have very different effects [52,53]. This highlights that DNA context affects epigenome editing in ways that are not currently understood.

Several studies have suggested that the ectopic insertion of an expanded CAG/CTG repeat in mice could induce changes in chromatin structure in the abutting sequences. An early example was the random insertion of arrays of transgenes, each carrying 192 CAGs, which led to the silencing of the transgenes independently of the site of genomic integration [19]. In addition, inserting a 40 kb human genomic region containing the *DMPK* gene along with an expansion of 600 CTGs [18], or a 13.5Kb region containing the human SCA7 gene with 92 CAGS [15] all led to changes in chromatin marks near the expansion. It has been unclear, however, whether the presence of endogenous sequence elements, like CpG islands [54] and CTCF binding sites [14,55], is necessary for this effect. Our data show that 91 CAGs, without the flanking sequences normally present at the *DMPK* gene from whence this repeat was cloned [30], does not lead to significant changes in the levels of H3ac in its vicinity. These data suggest that the flanking sequence elements may play important roles in the induction and/or maintenance of heterochromatic marks surrounding expanded CAG/CTG repeats.

We designed PInT specifically to isolate expanded repeats tracts from other potential locus-specific cis elements. This is helpful in finding factors that would affect instability and/or gene expression across several diseases. A potential application of PInT includes cloning in specific cis elements, including CTCF binding sites and CpG islands, next to the repeat tract and evaluate their effects on instability with or without targeting. In fact, PInT can be used to clone any sequence of interest near the targeting site and can be utilized for a wide array of applications, beyond the study of expanded CAG/CTG repeats.

PInT can be used to design peptides with enough activity to be useful in downstream epigenome editing applications. For instance, here we dissected the mechanisms of action of HDAC5 in silencing using mutants and truncations. We could quickly screen for domains and mutants that are effective in modulating gene expression. This is especially desirable in designing epigenome editing approaches with dCas9 fusions *in vivo*. A current limitation of the *S. pyogenes* Cas9 for *in vivo* applications is its large size, which is at the limit of what adeno-associated viral vectors can accommodate [56]. Even with the smaller orthologues, packaging a dCas9 fusion inside a gene delivery vector is a challenge, let alone encoding the sgRNA in the same vector. Therefore, being able to trim a chromatin modifier down to its smallest active peptide may help in optimizing downstream applications and translation.

In this study, we addressed a central question for both HDAC5 and DNMT1 and their involvement in CAG/CTG repeat instability. It has been unclear what the exact roles of these two enzymes might be in repeat instability. Specifically, whether they work by modifying the local chromatin structure or *in trans* has remained an outstanding issue. For DNMT1, it was speculated that increases in CpG methylation surrounding the repeat tract might facilitate repeat expansion [31]. Our data do not support such a model and rather point to an indirect role for DNMT1 in repeat instability, perhaps through changes in the transcriptome. For example, DNMT1 controls the expression of MLH1 [57], which has been shown to be important for repeat instability [58–61]. It should be pointed out that there remains the possibility that DNMT1 targeting did not lead to large enough changes in CpG methylation to affect repeat instability. The case of HDAC5 is possibly more complex as its partner, HDAC3, has been speculated to play a role in the deacetylation of MSH3 [33], which is a known modifier of repeat instability [33,59,70,71,62–69]. The results obtained with PInT are concordant with a role for HDAC5 *in trans*, which may be that it helps control the deacetylation of MSH3 before it binds to the repeat tract. These results highlight the usefulness of PInT in understanding the mechanism of repeat instability.

We found that targeting of PYL-HDAC3 increases gene expression slightly, independently of repeat size and in the presence of a small molecule inhibitor of its catalytic activity. Although this appears counterintuitive, several studies suggest that this is not unexpected. Specifically, HDAC3 has an essential role in gene expression during mouse development that is independent of its catalytic activity [72]. Moreover, HDAC3 binds more readily to genes that are highly expressed in both human and yeast cells [73,74]. The mechanism or function of HDACs binding to highly expressed genes are currently unknown.

The observation that expanded CAG/CTG repeats resist gene silencing is intriguing. This effect appears to be independent of which silencer is targeted as we have tried two, DNMT1 and HDAC5, which have very different modes of action. We have identified an HDAC3 inhibitor, RGFP966, that abolishes the difference in repeat size upon HDAC5 targeting without affecting the silencing activity. Importantly, we cannot currently rule out that RGFP966 may inhibit other HDACs that would be responsible for this effect. Epigenome editing, through targeting of PYL-HDAC5 or PYL-DNMT1, remained unaffected, with similar levels of deacetylation and DNA methylation levels regardless of repeat size. These results suggest that neither H3ac nor DNA methylation are good proxies for gene silencing. There are several steps towards gene silencing that could be differentially affected by the presence of a CAG/CTG repeat expansion. First there is transcription, but it is known to be impeded, at least *in vitro*, by the presence of a repeat tract [75]. This is counter to the effect on gene silencing that we observed here. Alternatively, splicing may be differentially regulated by both the expanded repeats and the targeted epigenome editing. Indeed, histone marks correlate with changes in splicing patterns [76] and expanded CAG/CTG repeats are known to affect splicing [30,77]. Moreover, both HDAC3 and HDAC5 interact with splicing factors [78]. Finally, we cannot rule out that mRNA or GFP stability may also contribute to the repeat-size-specific effect seen here, but the mechanism would have to be more convoluted. Ultimately, finding the HDAC3 target that mediates this effect will help understanding the mechanism of allele-specific gene silencing that we uncovered here.

The observations that expanded CAG/CTG repeats reduces the efficiency of gene silencing has implications in the design of epigenome editing approaches for expanded repeat disorders. We speculate that PInT may be adapted to screen for allele length-specific silencers, which may help design novel therapeutic options for expanded CAG/CTG repeat disorders.

## Materials and Methods

### Cell culture conditions and cell line construction

Most of the cell lines used, including all the parental lines, were genotyped by Microsynth, AG (Switzerland) and all confirmed to be HEK293.2sus. They were free of mycoplasma as assayed by the Mycoplasma check service of GATC Biotech. The cells were maintained at 37 °C with 5% CO_2_ in DMEM containing 10% FBS, penicillin, and streptomycin, as well as the appropriate selection markers at the following concentrations: 15 µg ml^-1^ blasticidin, 1µg ml^-1^ puromycin, 150µg ml^-1^ hygromycin, 400 µg ml^-1^ G418, and/or 400 µg ml^-1^ zeocin. Whereas FBS was used to maintain the cells, dialyzed calf serum was used at the same concentration for all the experiments presented here. The ABA concentration used was 500 µM, unless otherwise indicated. Doxycycline (dox) was used at a concentration of 2 µg ml^-1^ in all experiments. RGFP966 was used at a concentration of 10 µM. Notably, it is not possible to obtain several stable lines with the exact same repeat size as they are, by their nature, highly unstable. This is why we have lines with different repeat sizes. Furthermore, the sizes can change over time and upon thawing from the freezer.

A schematic of cell line construction and pedigree is found in Figure S1, and the lines are listed in Table S1. This table includes the plasmids made for cell line construction. The levels of the transgenes are found in Figure S3. The plasmids used for transient transfections are found in Table S2. For each cell line, single clones were isolated and tested for expression of ParB-ABI and PYL-fusions by western blotting using the protocol described before [11]. Briefly, whole cell extracts were obtained, and their protein content was quantified using the Pierce BCA Protein Assay Kit (ThermoScientific). Proteins were then run onto Tris-glycine 10% SDS PAGE gels before being transferred onto nitrocellulose membrane (Axonlab). The membranes were blocked using the Blocking Buffer for Fluorescent Western Blotting (Rockland), and primary antibodies were added overnight. Membranes were then washed followed by the addition of the secondary antibody (diluted 1 to 2000). The fluorescent signal was detected using an Odyssey Imaging System (Li-CoR). All antibodies used are found in Table S3. To assess repeat sizes, we amplified the repeat tracts using oVIN-0459 and oVIN-0460 with the UNG and dUTP-containing PCR as described [79] and then Sanger-sequenced by Microsynth AG (Switzerland). The sequences of all the primers used in this study are found in Table S4.

The ParB-INT sequence system used here is the c2 version described previously [23], except that the ParB protein was codon-optimized for expression in human cells. It is also called ANCHOR1 and is distributed by NeoVirTech. ParB-ABI (pBY-008), PYL (pAB-NEO-PYL), PYL-HDAC5 (pAB(EXPR-PYL-HDAC5-NEO)) and PYL-HDAC3 (pAB(EXPR-PYL-HDAC3-NEO)) constructs were randomly inserted and single clones were then isolated (Table S1). GFP-reporter cassettes were inserted using Flp-mediated recombination according to the manufacturer’s instruction (Thermo Scientific). Single colonies were picked and screened for zeocin sensitivity to ensure that the insertion site was correct.

### Targeting assays

Detailed protocols of the assay and culture conditions can be found in [80]. For targeting assays involving transient transfections, cells were plated onto poly-D-lysine-coated 12-well plates at a density of 6×10^5^ cells per well and transfected using 1 µg of DNA per well and Lipofectamine 2000 or Lipofectamine 3000 (Thermofisher Scientific). 6 hours after transfection, the medium was replaced with one containing dox and ABA or DMSO. 48h after the transfection, the cells were split, and fresh medium with dox and ABA or DMSO was replenished. On the fifth day, samples were detached from the plate with PBS + 1 mM EDTA for flow cytometry analysis.

In the case of the stable cell lines, cells were seeded at a density of 4×10^5^ per well in 12-well plates. The media included dox and ABA or DMSO. The medium was changed 48 hours later and left to grow for another 48 hours. The cells were then resuspended in 500µl PBS + 1 mM EDTA for flow cytometry analysis.

### Flow cytometry

We used an Accuri C6 flow cytometer from BD and measured the fluorescence in at least 12 500 cells for each treatment. The raw data was exported as FCS files and analysed using FlowJo version 10.0.8r1. A full protocol is available here [80].

### Chromatin immunoprecipitation

For chromatin immunoprecipitation, the cells were treated as for the targeting experiments except that we used 10 cm dishes and 4×10^6^ cells. After 96 hours of incubation, paraformaldehyde was added to the medium to a final concentration of 1% and the cells were incubated for 10 minutes at room temperature. The samples were then quenched with 0.125 M PBS-glycine for 5 minutes at room temperature. Samples were then centrifuged, the supernatant was discarded, and the cell pellets were washed with ice-cold PBS twice. The samples were split into 10^7^ cell aliquots and either used immediately or stored at -75 °C for later use. Sonication was done using a Bioruptor for 25 to 30 min. DNA shearing was visualized by agarose gel electrophoresis after crosslink reversal and RNase treatment. 20% of sonicated supernatant was used per IP, with 3 μg anti-FLAG (M2, Sigma), anti-PAN acetylated H3 (Merck), or anti-IgG (3E8, Santa Cruz Biotechnology) on Protein G Sepharose 4 Fast Flow beads (GE healthcare). The samples were incubated at 4°C overnight and then washed with progressively more stringent conditions. After the IP, the samples were de-crosslinked and purified using a QIAquick PCR purification kit (Qiagen) and analysed using a qPCR.

### Quantitative PCR

Quantitative PCR was performed with the FastStart Universal SYBR Green Master Mix (Roche) using a 7900HT Fast Real-Time PCR System in a 384-Well Block Module (Applied Biosystems™). Primers used to detect enrichment at the INT sequence and at *ACTA1* gene are listed in Table S4. Ct values were analysed using the SDS Software v2.4. The percentage of input reported was obtained by dividing the amount of precipitated DNA for the locus of interest by the amount in the input samples multiplied by 100%.

### Small pool PCR

Small pool PCR experiments were performed on DNA isolated from 91-Y, 59B-Y-HDAC5, and 89B-Y-DNMT1 cells grown with ABA or DMSO only for 30 days in the presence of the appropriate selection markers. The SP-PCR protocol used is described in [79]. We used primers oVIN-460 and oVIN-1425 (Table S5) to amplify the repeat tract, ran the products on a TAE agarose gel and alkaline transferred it on positively charged nylon membrane (MegaProbe). The membranes were then probed with oVIN-100 (5’-CAGCAGCAGCAGCAGCAGCAGCAGCAGCAG) that was end-labelled with ^32^P and exposed to a phosphoscreen and scanned with a Typhoon scanner. To quantify the instability, membranes were blinded, and a different lab member drew lines at the top and bottom of the most common band and then counted individual alleles that fell outside of these lines. We estimated the total number of alleles amplified using a Poisson distribution based on the number of empty wells on our membranes as described [31]. We have noted that cell lines with repeats that are mildly expanded (e.g., 59 CAGs) have fewer contractions than longer ones. This is consistent with studies in the context of DM1 and HD [81], albeit the size threshold for seeing more contractions may be shorter in HEK293-derived cells than in mice.

### Bisulfite sequencing

Bisulfite conversion was done using the EZ DNA Methylation kit from Zymo Research as described before [20]. We converted 200ng of DNA at 50°C for 12 hours from each cell line after 30 days of culturing with ABA. We used primer oVIN-2209 and oVIN-2211 to amplify the converted DNA (Table S5). The products were then purified using the NucleoSpin PCR Clean-up kit (Macherey-Nagel). We then performed 2×250 bp paired-end MiSeq sequencing (Illumina).

The sequencing primers are found in Table S5. We processed the reads with TrimGalore (github.com/FelixKrueger/TrimGalore) using -q 20 --length 20 –paired. We aligned the reads using QuasR [82] to the GFP transgene sequence. We extracted the methylation levels for each CpG in the amplicon with the qMeth() function in QuasR. We calculated the CpG methylation frequencies by dividing the frequency of methylated CpGs by the total number of CpG and expressed it as a percentage.

### Statistics

We determined statistical significance in the targeting and ChIP experiments using a two-tailed one-way ANOVA. For small-pool PCR we used a χ^2^ test with two degrees of freedom using three categories: expansions, contractions, and no change. We used a Fisher’s exact test in the case of the 59B-Y-HDAC5 lines because we found no contractions. All the statistical analyses were done using R studio version 3.4.0. We concluded that there was a significant difference when P < 0.05.

## Acknowledgements

We thank John H. Wilson and Kerstin Bystricky for sharing reagents, as well as Fisun Hamaratoglu, Helder C. Ferreira, Ana C. Marques, Johanna E. Ainsworth, Nastassia Gobet, Emma Randall, and Marcela Buricova for critical reading of the manuscript. Florence Gidney blindly counted the unstable alleles. This work was supported by the Swiss National Science Foundation Professorship to V.D. (#172936). Work in V.D.’s lab is also supported by the UK Dementia Research Institute, which receives its funding from DRI Ltd, funded by the UK Medical Research Council, Alzheimer’s Society and Alzheimer’s Research UK.

## Author contributions

BY performed the experiments unless stated otherwise. ACB performed all the experiments for Fig. 6. The small pool PCRs in Fig. 2,3, and 4 and in Table 1 were done by ORL. GARB performed the bisulfite sequencing in Fig. 4; TB analysed these data. ACB and LA helped BY generating the cell lines. BY, ACB, and VD designed the experiments. CC performed the statistical analyses. BY and VD wrote the paper and prepared the figures.

## Funding

This work was funded by SNSF professorship grants #144789 and #172936 to V.D. VD is also supported by the UK Dementia Research Institute, which receives its funding from DRI Ltd, funded by the UK Medical Research Council, Alzheimer’s Society and Alzheimer’s Research UK.

## Data availability statement

The datasets, cell lines, and plasmids generated and analysed in the current study are available from the corresponding author. Note that to obtain some of the plasmids you will also need the permission of NeoVirTech, which owns the rights to the ANCHOR technology.

## Supplemental Materials

**Table S1:**
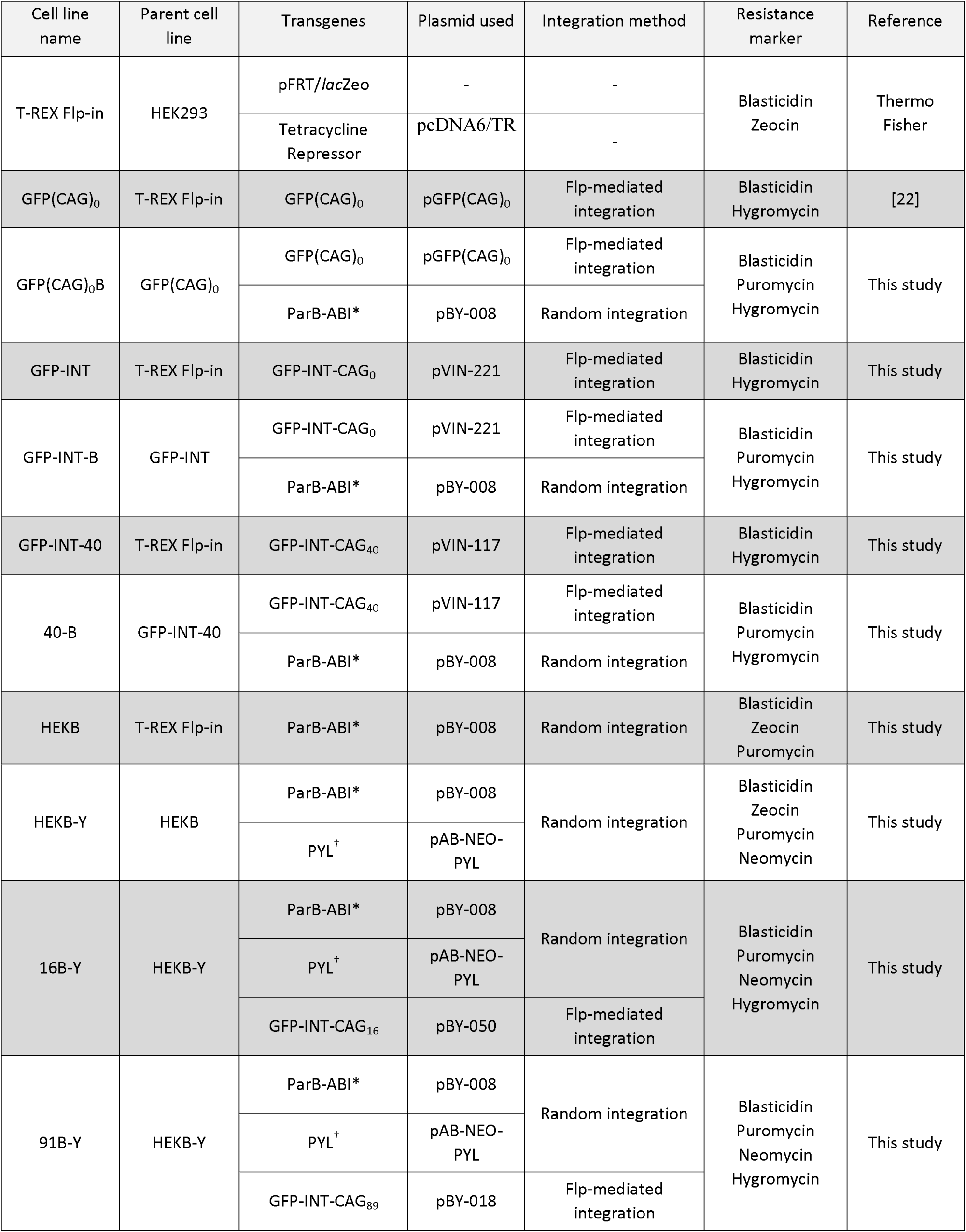
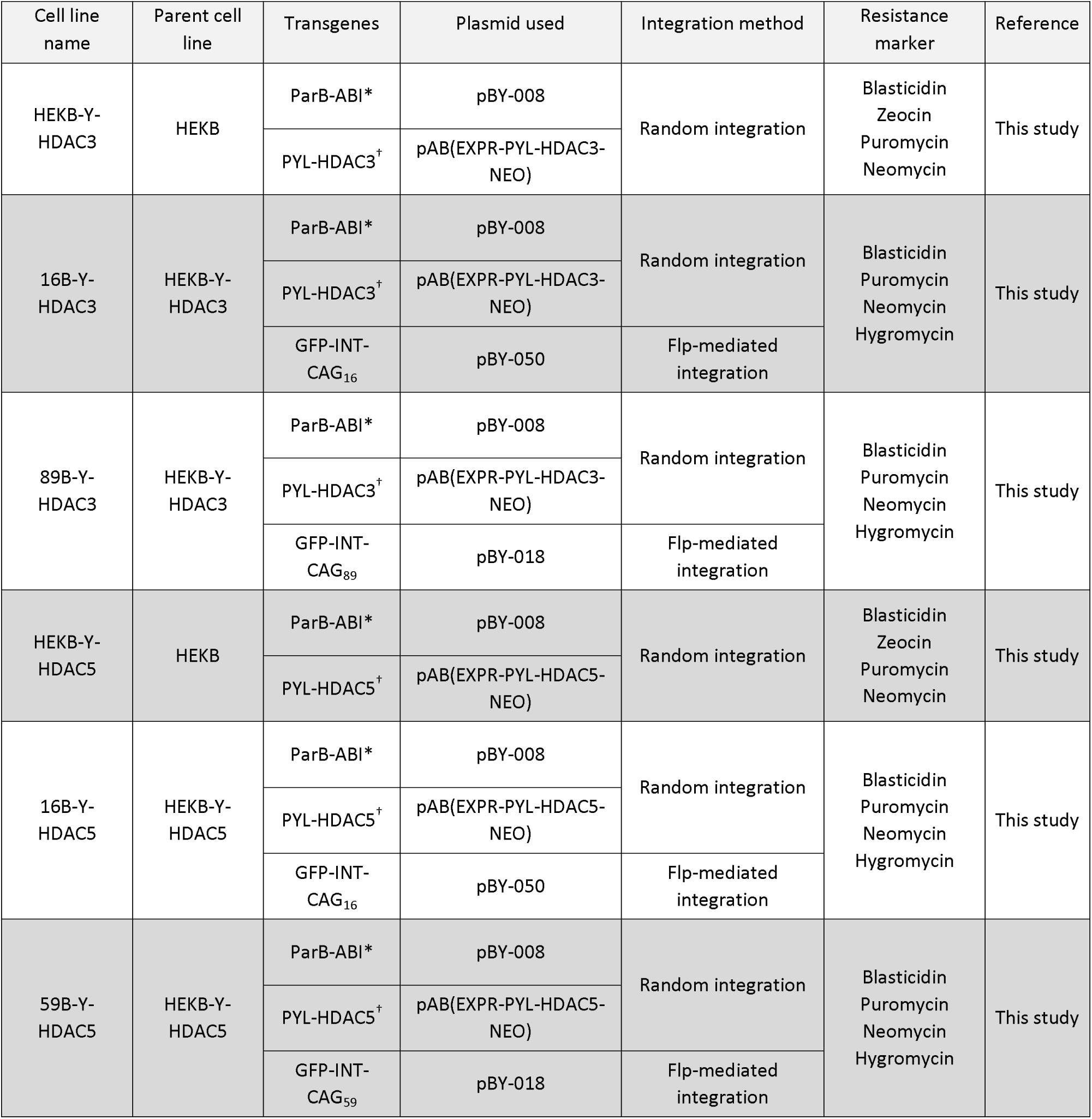
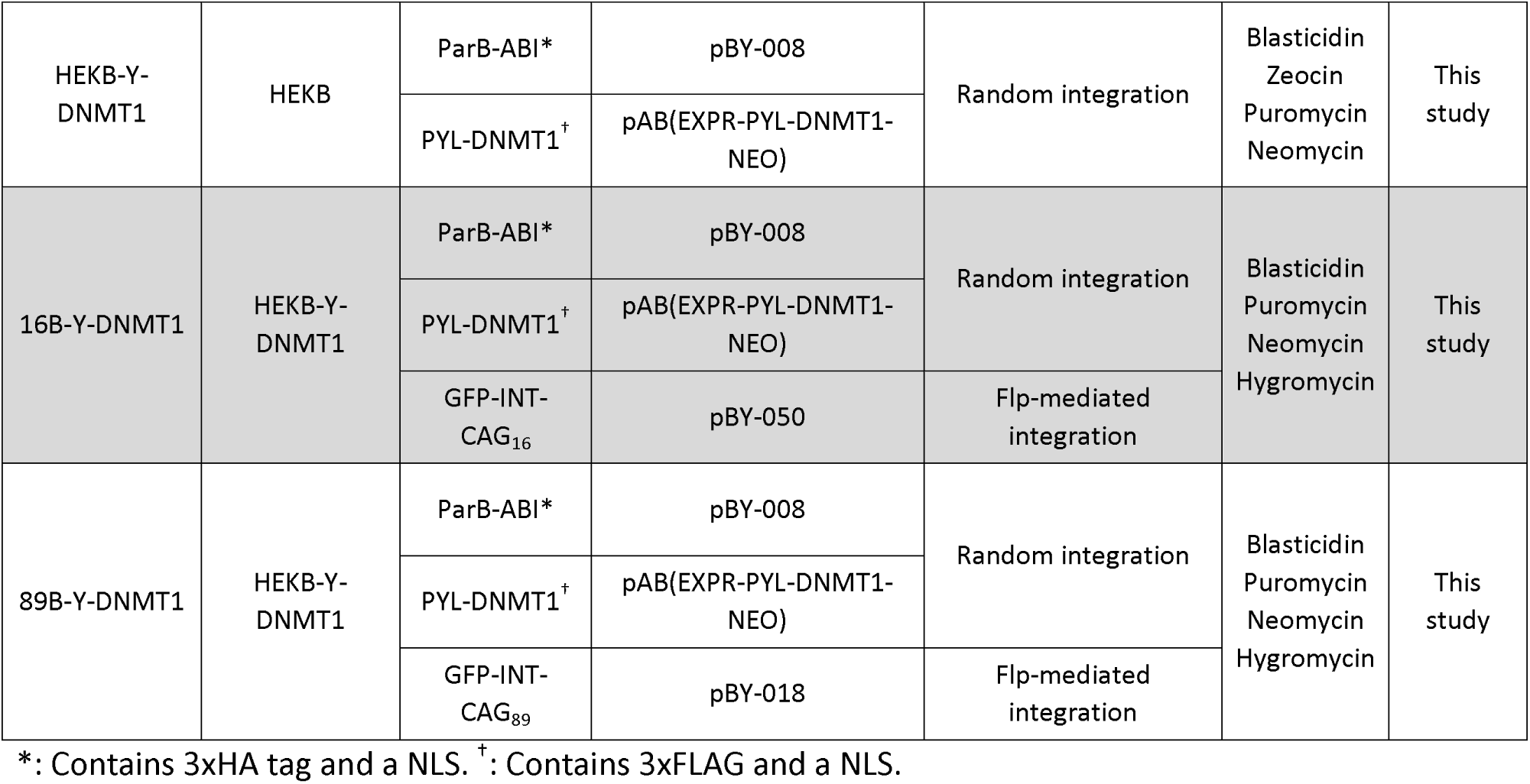
Cell lines.

**Table S2:** Plasmids used for transient transfection experiments.

| Name | Description | Source |
| --- | --- | --- |
| pBY-008 | Expresses ParB-ABI with 3xHA and a SV40 NLS | This study |
| pBY-022 | Expresses PYL fused to 3xFLAG and a SV40 NLS. Also serves as a destination vector for making fusions | This study |
| pBY-006 | Expresses PYL-HDAC3 with 3xFLAG and a SV40 NLS | This study |
| pBY-017 | Expresses PYL-HDAC5 with 3xFLAG and a SV40 NLS | This study |
| pAB-HDAC5(mut-H-A) | Expresses PYL-HDAC5 H1006A with 3xFLAG and a SV40 NLS | This study |
| pAB-HDAC5(mut-H-Y) | Expresses PYL-HDAC5 H1006Y with 3xFLAG and a SV40 NLS | This study |
| pAB(EXPR-HDAC5-trunc1) | Expresses PYL-HDAC5 aa 1-275 with 3xFLAG and a SV40 NLS | This study |
| pAB-EXPR(PYL-cat_dom_HDAC5) | Expresses the PYL-HDAC5 catalytic domain with 3xFLAG and a SV40 NLS | This study |
| pAB-EXPR(PYL-Nterm_dom_HDAC5) | Expresses the PYL-HDAC5 N terminal domain fused to 3xFLAG and a SV40 NLS | This study |
| pAB(EXPR-PYL-DNMT1-NEO) | Expresses PYL-DNMT1 with 3xFLAG and a SV40 NLS | This study |

**Table S3:** Antibodies.

| Epitope | Company | Catalog number | Dilution | Assay |
| --- | --- | --- | --- | --- |
| FLAG | Sigma-Aldrich | F1804-5MG | 3 µg per IP | ChIP |
|  |  |  | 1:1000 | WB |
| HA | Sigma-Aldrich | 12158167001 | 1:2000 | WB |
| IgG | Santa Cruz Biotechnology | sc-69786 | 3 µg per IP | ChIP |
| Pan-acetylation of H3 | Merck | #06-599 | 3 µg per IP | ChIP |
| Histone H3 | Abcam | ab1791 | 3 µg per IP | ChIP |
| Actin | Sigma-Aldrich | A2066-.2ML | 1:2000 | WB |

**Table S4:**
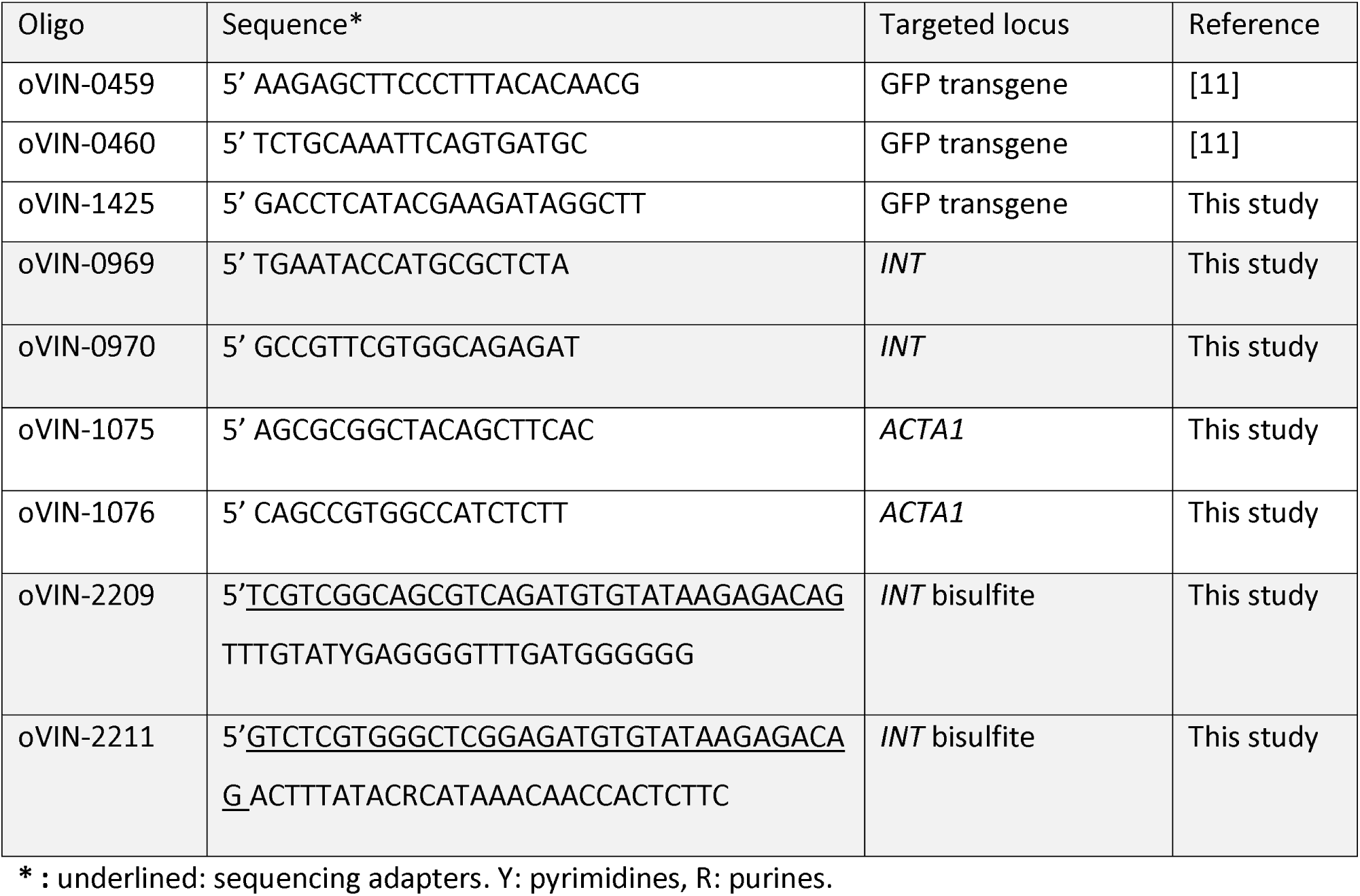
Primers using in this study.

**Fig. S1.**
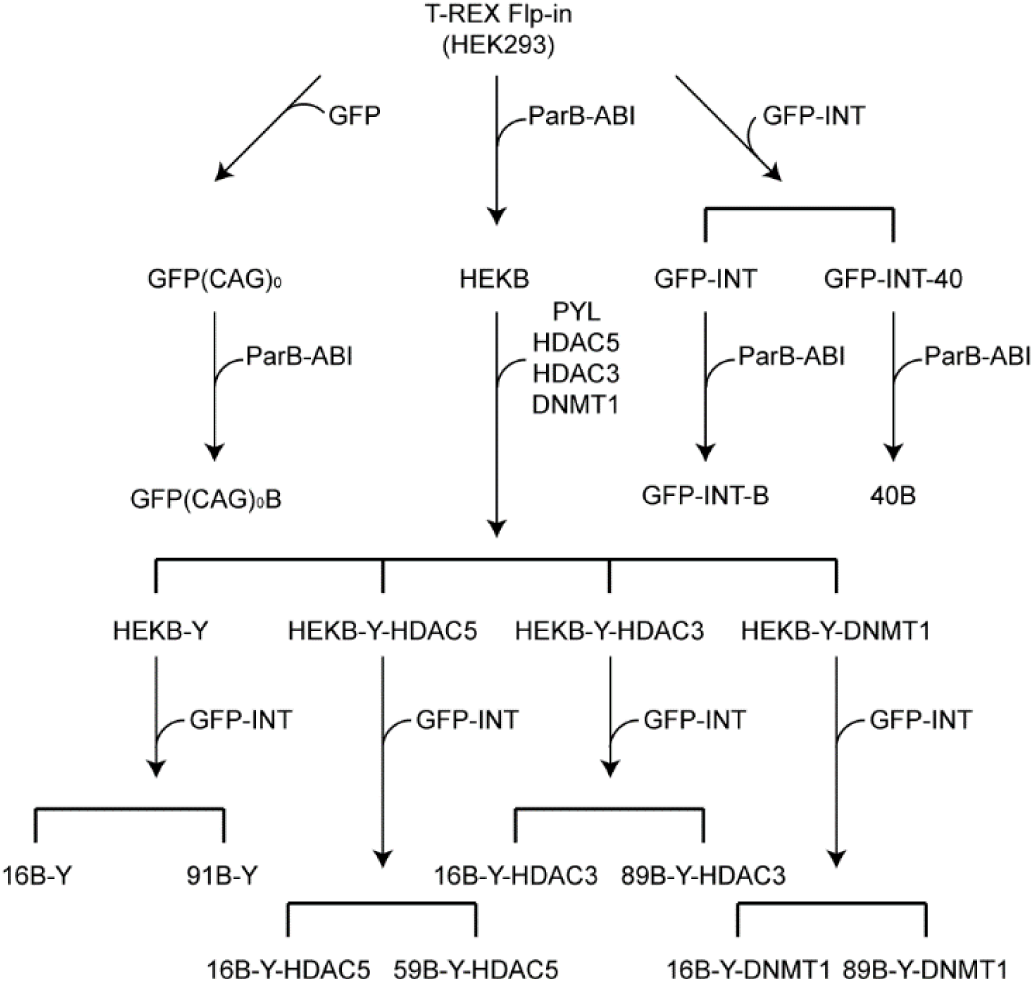
Parentage of the cell lines described in this study. All cell lines are clonal. Details of each plasmid integrated, the methods of doing so, and the details of the cell lines are found in Tables S1.

**Fig. S2.**
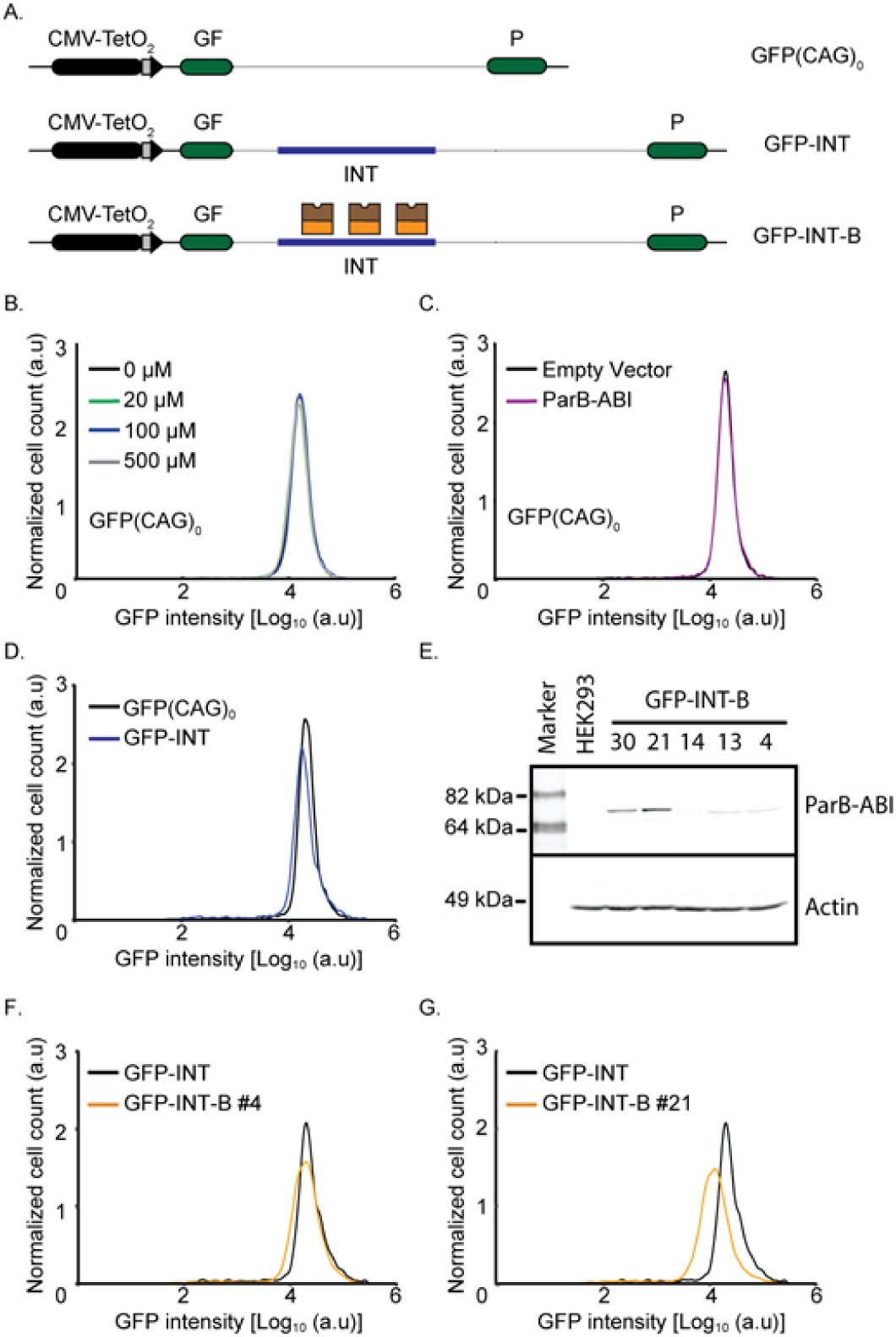
Effect of components of PInT on GFP expression. A) Cartoon of cell lines used. B) Representative flow cytometry profiles of GFP(CAG)_0_ cells treated with increasing concentrations of ABA dissolved into the same volume of DMSO. C) Representative flow cytometry profiles of GFP(CAG)_0_ cell lines transfected with a plasmid expressing ParB-ABI or an empty vector. D) Comparison of GFP expression between GFP(CAG)_0_ and GFP-INT cells. E) Western blots (against HA) of GFP-INT-B clones showing varying amounts of ParB-ABI in the different clones. F) Flow cytometry profiles of GFP-INT-B clone #4, which expresses low levels of ParB-ABI. G) Representative flow cytometry profile of GFP-INT-B clone #21 expressing a larger amount of ParB-ABI. Note that the GFP-INT parent profile is the same in panels F and G because the GFP expression of both clones was done on the same day using the same parent cell line as control.

**Fig. S3.**
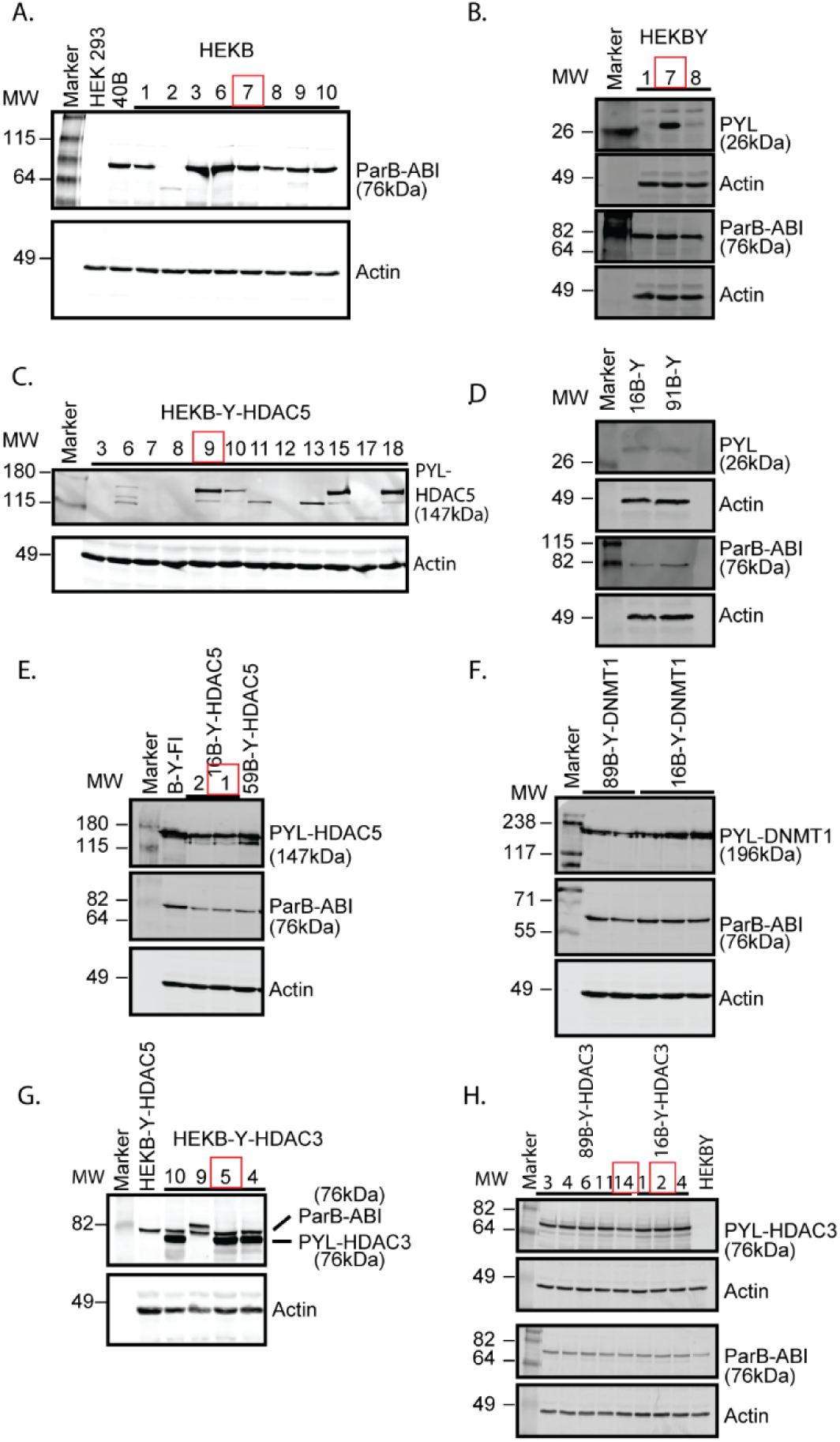
Cell line making and characterization. The levels of ParB-ABI and PYL-fusions by western blotting using antibodies against HA and FLAG, respectively. The red boxes identify the clones that were used subsequently. A) ParB-ABI levels in HEKB clones. B) PYL and ParB-ABI levels in HEKBY cells. C) PYL-HDAC5 in HEKB-Y-HDAC5 cells. D) PYL and ParB-ABI levels in 16B-Y and 91B-Y cells E) PYL-HDAC5 and ParB-ABI levels in 16B-Y-HDAC5 and 59B-Y-HDAC5 cells. F) PYL-DNMT1 and ParB-ABI levels in 89B-Y-DNMT1 and 16B-Y-DNMT1 cells. G) PYL-HDAC3 and ParB-ABI levels in HEKBYH3 cells. H) PYL-HDAC3 and ParB-ABI levels in 16B-Y-HDAC3 and 89B-Y-HDAC3 cells. The MW marker used was the BenchMark™ Pre-stained Protein Standard, except in (F), where the HiMark™ Pre-stained Protein Standard was used.

**Fig. S4.**
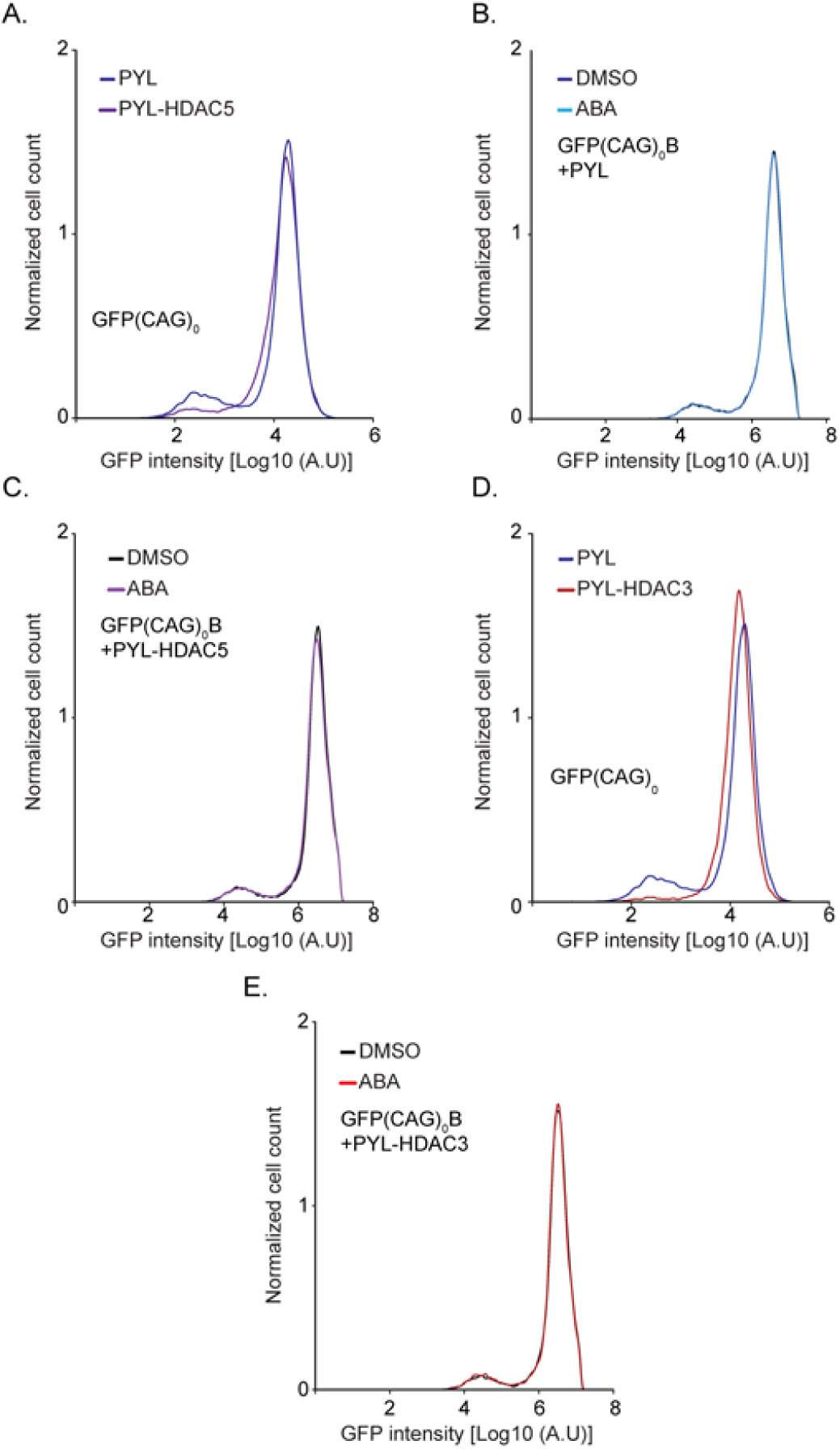
The functionality of PYL fusions in GFP-INT-B cells and GFP(CAG)_0_B cells. Representative flow cytometry profiles after transient overexpression of PYL and PYL-HDAC5 in GFP-INT-B cells (A), or of PYL and PYL-HDAC3 in GFP-INT-B cells (B). C-E) Representative flow cytometry profiles of GFP(CAG)_0_B cells transiently transfected with PYL (C), PYL-HDAC5 (D), or PYL-HDAC3 (E) and incubated with DMSO or ABA.

**Fig. S5.**
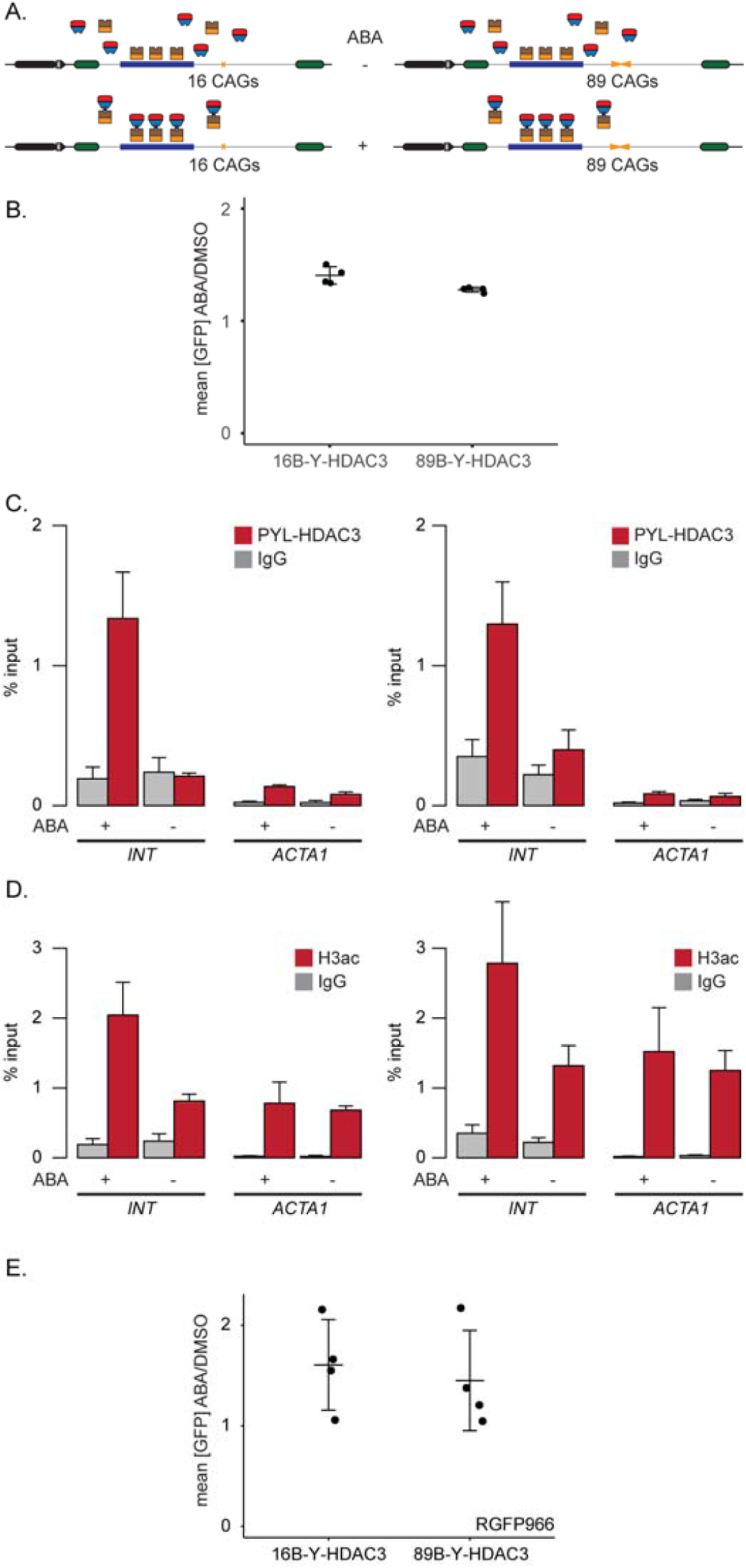
Targeting of PYL-HDAC3 increases GFP expression independently of its catalytic activity. A) Schematic representation of the nB-Y-HDAC3 cells. B) Quantification of GFP expression in nB-Y-HDAC3 cells with ABA or DMSO (16B-Y-HDAC3: N=4; 89B-Y-HDAC3: N=4). The error bars are the standard error around the indicated mean. C) ChIP-qPCR experiments using an antibody against PYL-HDAC3 fusion (FLAG) at the INT and ACTA1 loci in the presence of ABA or DMSO. Left, 16B-Y-HDAC3 cells (N=4), Right, 89B-Y-HDAC3 cells (N=4). The error bars are the standard error. D) ChIP-qPCR experiments using an antibody against pan-acetylated H3 (H3ac) at the INT and ACTA1 loci in the presence of ABA or DMSO. Left, 16B-Y-HDAC3 cells (N=4), Right, 89B-Y-HDAC3 cells (N=4). The error bars are the standard error. E) Quantification of GFP expression in nBYH3 cells with ABA or DMSO and treated with RGFP966 (16B-Y-HDAC3: N=4; 89B-Y-HDAC3: N=4). The error bars are the standard error around the indicated mean.

**Fig. S6:**
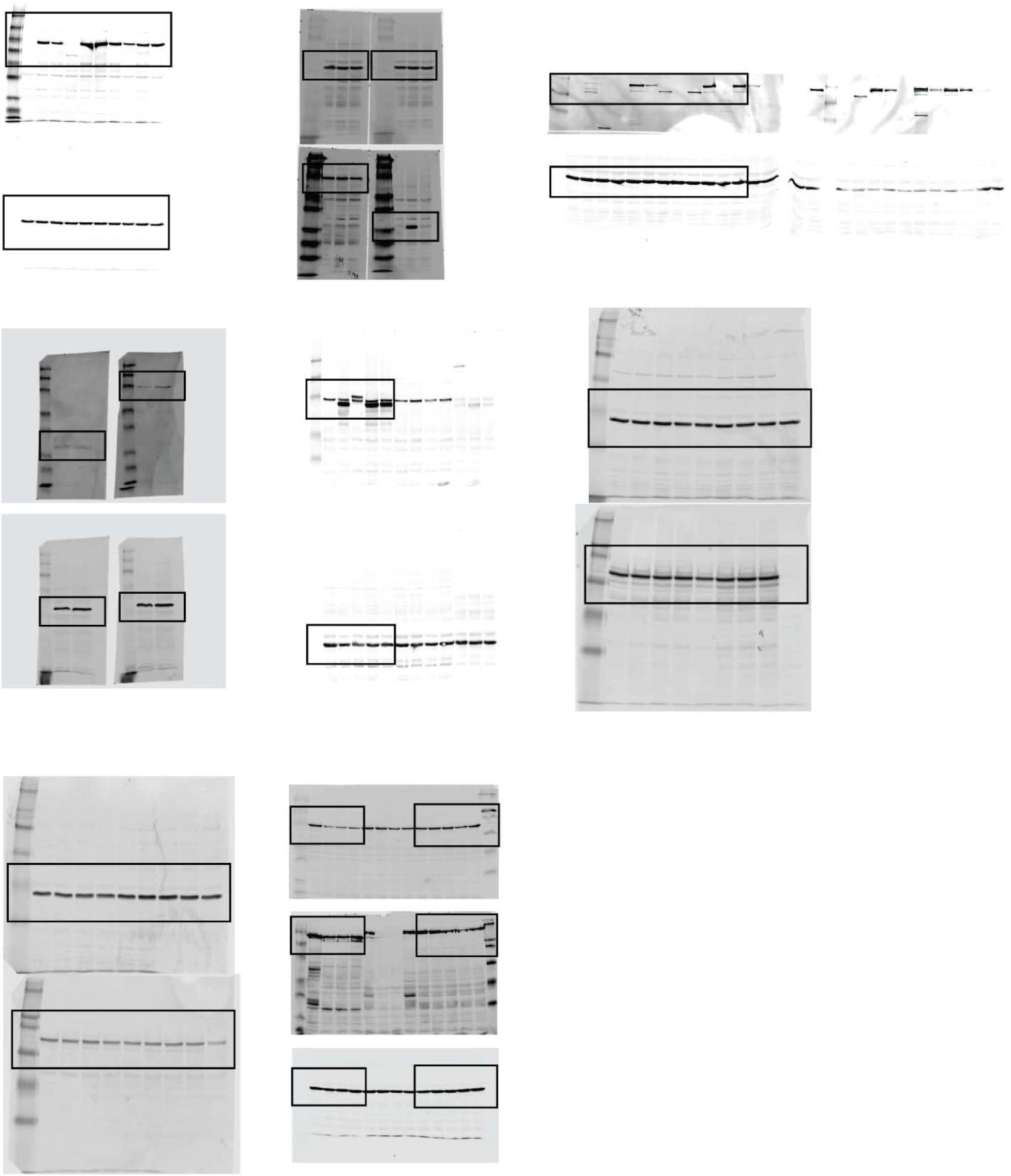
Unaltered full western blots. Black boxes indicate where the blots were cropped.

**Fig. S7:**
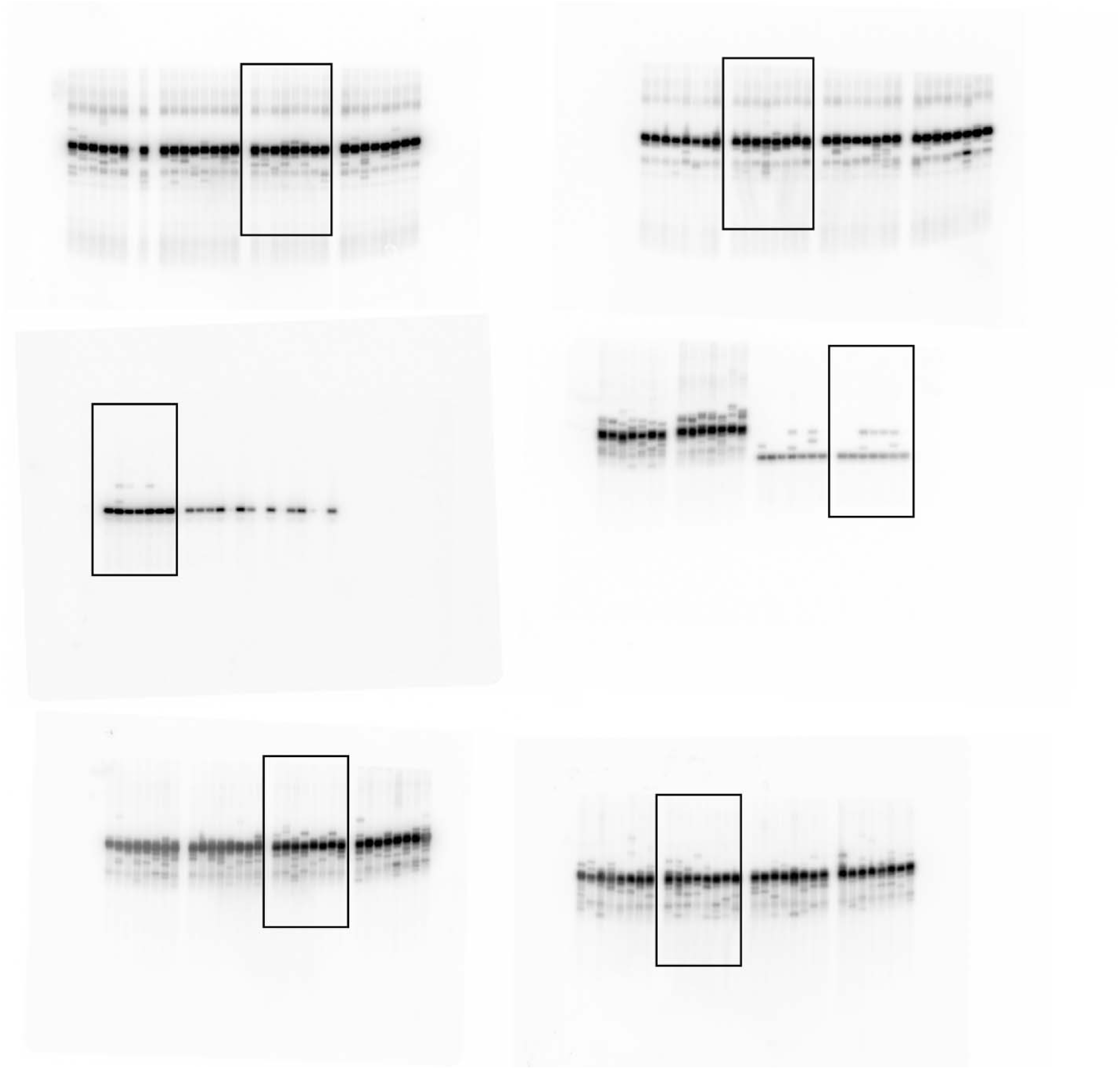
Unaltered full small-pool PCR blots. Black boxes indicate where the blots were cropped.

## Notes

### Competing Interest Statement

The authors have declared no competing interest.

